# Reactivation of single-episode pain patterns in the hippocampus and decision making

**DOI:** 10.1101/2020.05.29.123893

**Authors:** G. Elliott Wimmer, Christian Büchel

## Abstract

Aversive and rewarding experiences can exert a strong influence on subsequent behavior. While decisions are often supported by the value of single past episodes, most research has focused on the role of well-learned value associations. Recent studies have begun to investigate the influence of reward-associated episodes, but it is unclear if these results generalize to negative experiences such as pain. To investigate whether and how the value of previous aversive experiences modulates behavior and brain activity, in our experiments female and male human participants experienced episodes of high or low pain in conjunction with incidental, trial-unique neutral pictures. In an incentive-compatible surprise test phase, we found that participants avoided pain-paired objects. In a separate fMRI experiment, at test, participants exhibited significant pain value memory. Neurally, when participants were re-exposed to pain-paired objects, we found no evidence for reactivation of pain-related patterns in pain-responsive regions such as the anterior insula. Critically, however, we found significant reactivation of pain-related patterns of activity in the hippocampus, such that activity significantly discriminated high versus low pain episodes. Further, stronger reactivation in the anterior hippocampus was related to improved pain value memory performance. Our results demonstrate that single incidental aversive experiences can build memories that affect decision making and that this influence may be supported by the hippocampus.

**Significance Statement:** Aversive and rewarding experiences can exert a strong influence on our subsequent behavior. While decisions are often supported by single past negative or positive episodes, most research has focused on the role of well-learned value associations. In experiments using aversive heat pain in conjunction with incidental objects, we found that participant’s choices were biased by the level of pain associated with the objects. Further, when participants saw the objects again, pain-related neural patterns in the hippocampus were re-expressed and this was related to pain value memory performance. These results suggest a mechanism by which even single negative experiences can guide our later decisions.

## Introduction

Our decisions are oriented toward seeking out rewarding experiences and, conversely, avoiding negative experiences. When faced with a choice of how to get to a restaurant, we may use different kinds of memories to avoid a negative experience: we may be biased against taking the bus because it is always delayed, or against taking a particular subway route because the on the last ride, the train was unbearably hot. Research on learning and decision making has predominantly focused on the influence of well-learned values on choice (Daw and Doya, 2006; Schultz, 2006; Rangel et al., 2008). However, our behavior is often influenced by single past experiences. Rapidly learning to avoid negative events from even a single exposure can be critical for survival, yet we know surprisingly little about the neural mechanisms that support the use of such memories in value-based decision making (Wimmer and Buchel, 2016).

In the last few years, research in decision making has benefitted from becoming more integrated with research in memory, building on proposals that value-based choice can be supported by a mechanism that samples representations stored in memory (Hertwig et al., 2004; Stewart et al., 2006; Weber and Johnson, 2006; Biele et al., 2009; Gluth et al., 2015; Shadlen and Shohamy, 2016). Importantly, for memories to guide value-based choices, those memories often need to be combined with the positive or negative value of the original experience. Early studies in behavioral economics demonstrated that participants can compare the aversive value of two past episodes, such as different experiences of unpleasant cold water or aversive film clips (Fredrickson and Kahneman, 1993; Kahneman et al., 1993; Redelmeier and Kahneman, 1996). Building on this, recent studies of decision making in the reward domain have shown an influence of single past episodes on decision making (Duncan and Shohamy, 2016; Murty et al., 2016; Wimmer and Buchel, 2016; Bornstein et al., 2017; Bornstein and Norman, 2017).

The hippocampus is critical for episodic memory, and relational memory more generally (Eichenbaum and Cohen, 2001; Davachi, 2006), suggesting that it could also play a critical role in associating episodes with value. Thus far, however, no studies have demonstrated a role for the hippocampus in the implicit learning of values from episodes. Activation in the hippocampus has been shown to correlate with the value of stimuli and snack foods (Lebreton et al., 2009; Gluth et al., 2015). Studies have also reported that the hippocampus is associated with decision making processes for well-learned values such as snack foods, potentially implementing a memory sampling mechanism (Gluth et al., 2015; Bakkour et al., 2019). In our previous study of incidental episodic reward associations, using multivariate techniques we found reactivation of reward-related regions but no effects in the hippocampus (Wimmer and Buchel, 2016). However, the previous study employed brief experiences; by increasing episode length and separation (Ezzyat and Davachi, 2011), it may be possible to better test a role of the hippocampus in value memory. Interestingly, relational memory linking an element with value may even be unrelated to traditional measures of episodic memory (e.g. Wimmer and Shohamy, 2012; Wimmer and Buchel, 2016).

The anterior hippocampus in particular may play an important role in encoding associations between episodes and value, given research demonstrating a central role for the anterior hippocampus in anxiety (Adhikari et al., 2010; Fanselow and Dong, 2010; Bach et al., 2014) as well as in memory integration and generalization (Poppenk et al., 2013; Schlichting et al., 2015; Brunec et al., 2018). Particularly for negative experiences, understanding the role of the hippocampus in value memory may be important for the understanding mood disorders and post-traumatic stress disorder (Hamilton and Gotlib, 2008; Brewin et al., 2010; Shin and Liberzon, 2010).

In contrast, the gradual learning of stimulus-value associations over multiple experiences is known to involve systems including the dopaminergic midbrain, striatum, insula, and amygdala (Schultz et al., 1997; LeDoux, 2000; Seymour and al., 2004; Schiller et al., 2008). In the case of learning from aversive stimuli such as heat, a network of pain-responsive regions including the insula and secondary somatosensory cortex is an additional likely substrate for memory for the value of pain (Seymour et al., 2004; Apkarian et al., 2005; Tracey and Mantyh, 2007; Roy et al., 2014; Horing et al., 2019).

In the following experiments, we investigated whether single aversive episodes influence memory-based decision making and whether such an influence is supported by reactivation of distributed patterns of pain-related activity in the hippocampus and pain-responsive regions. During the incidental learning phase, neutral objects were presented once, incidentally paired with high or low pain (**Fig. 1a**). A surprise choice phase or a pain value memory test phase followed (**Fig. 1b-c**). By training a multivariate classifier on initial pain experience, at re-exposure, we could then test for reactivation of pain-related patterns and whether these effects were related to value memory performance.

**Figure 1.**
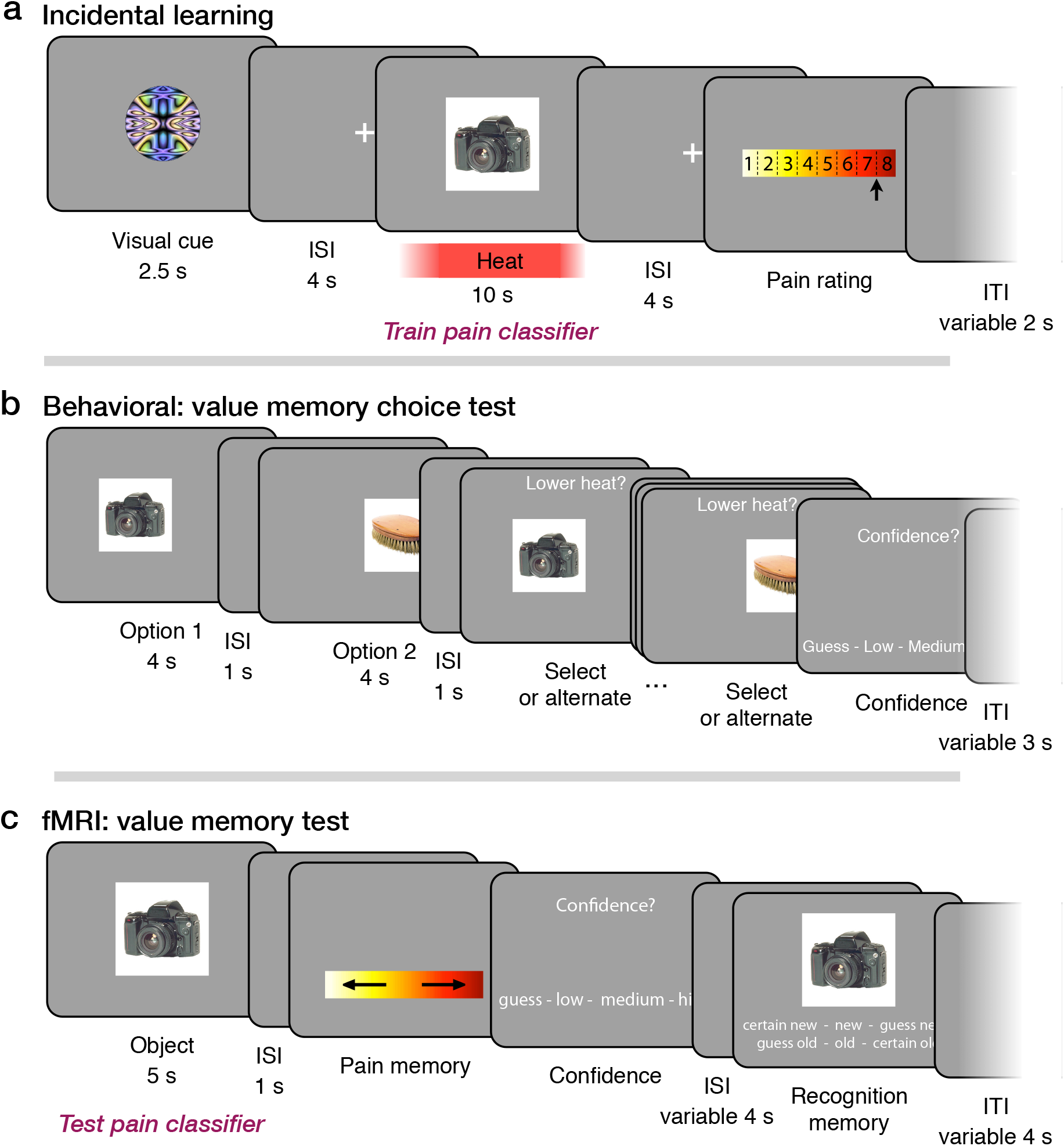
Pain value memory experimental design. ***a***, In the incidental learning phase, participants experienced high or low heat pain while being exposed to incidental trial-unique object pictures. Participants then rated their experienced level of pain. This phase was scanned in the fMRI study. ***b***, Value memory choice phase in the behavioral experiment. Each trial presented two objects from the incidental learning phase in sequence. Participants then alternated between objects to select the object that they thought had been associated with lower heat pain. ***c***, Value memory test phase in the fMRI experiment. Each trial presented a single object and participants responded with whether the object was paired with high or low heat pain and then rated their confidence in this response. Finally, participants then rated their recognition strength on a 6-point new-to-old scale.

## Materials and Methods

### Participants

A total of 26 subjects participated in the behavioral choice experiment. Participants were right-handed fluent German speakers with no self-reported neurological or psychiatric disorders. Data from two participants were excluded due to technical problems with the thermode and data from three additional participants were excluded due to errors in response recording, leaving 21 participants (13 female; mean age 25.1 years; range 18-42). A total of 31 subjects participated in the fMRI experiment. Participants were right-handed fluent German speakers with no self-reported neurological or psychiatric disorders and normal or corrected-to-normal vision. Data from two participants were excluded due to technical problems with the thermode, leaving 29 participants (15 female; mean age 26.0 years; range 20-33 years). In one participant, pain memory confidence ratings and memory recognition strength in the immediate test session were not recorded due to a technical error. The Ethics committee of the Medical Chamber Hamburg approved the study and all participants gave written consent.

### Experimental design

The experiments were designed to allow an investigation of the cognitive and neural mechanisms that support memory for aversive experiences, which is the focus of the current report. Secondarily, and separately, the experiment allows for an investigation of the behavioral and neural correlates of pain modulation of short term and very long-term recognition memory. For the latter question, a subset of participants returned one year later to assess whether the maintenance of recognition memory was modulated by pain and neural activity during the fMRI session; these results will be published separately (Wimmer and Buchel, 2015).

As an overview, the behavioral and fMRI experiments each started with a heat calibration phase. This was followed by the incidental learning phase (which was scanned in the fMRI experiment), where an abstract cue probabilistically associated with high or low heat were followed by the presentation of a trial-unique object in conjunction with high or low heat pain. A test phase followed that measured whether pain value memory could support rewarded choice of a low pain-associated object over a high pain-associated object (in the behavioral experiment) and single-item pain value memory (in the fMRI experiment).

The test phase was designed to be as sensitive as possible to behavioral signatures of pain value memory established via single episodes. Thus, we explicitly instruct participants to retrieve pain value associations. Such instruction differs from decisions about well-learned value associations, which can frame instructions in terms of preference. However, our use of many diverse stimuli prevents the use of such a general preference question: participants are bound to have idiosyncratic and widely varying preferences for the object pictures themselvesThus, if a preference instruction had been used, these idiosyncratic preferences would be likely to dominate behavioral and neural responses at test phase re-exposure. Further, due to the requirement for novel experiences, a pre-rating phase to collect baseline object ratings was not possible.

The test phase in the fMRI experiment allowed for the investigation of the critical question of whether pain-related patterns of activity were re-activated upon re-exposure to objects and whether individual differences in reactivation related to individual differences in value memory performance. Here, we employed multivariate methods, which are powerful and highly sensitive tools for investigating neural representations in memory and of pain experience (Poldrack, 2011; Rissman and Wagner, 2012; Wager et al., 2013).

### Heat calibration

Before the incidental learning phase, heat levels were calibrated for each participant to achieve the same subjective high and low aversive pain experience across participants. Thermal stimulation was delivered via an MRI compatible 3 × 3 cm Peltier thermode (MSA; Somedic, Sweden) applied to the inner left forearm. During the visual presentation of a white square, heat was applied for 10 s. Pain ratings were recorded with a 1-8 rating scale with 0.5-point increments, superimposed on a yellow-to-red gradient (as depicted in Fig. 1a). Participants moved an arrow cursor from the initial mid-point starting location using left and right key-presses and confirmed the rating by pressing the spacebar. A rating of ‘8’ corresponded to the highest level of heat pain a participant could endure multiple times. If the level of pain was intolerable, participants moved the rating past the ‘8’ end of the scale, at which point a ‘9’ appeared on the screen. Participants rated the pain associated with a pseudo-random list of 10 different temperatures ranging from 39.5 to 49.5°C. A linear interpolation algorithm then selected a low temperature estimated to yield a ‘2’ rating and a high temperature estimated to yield a ‘7.5’ rating.

To ensure no damage to participants’ skin due to the administered heat stimulation, the maximum temperature allowed in the experiment was 50.5 °C. Further, as described in detail below, if participants at any point entered a ‘9’ rating during the experiment, the high temperature was subsequently decreased by 0.8 °C.

### Procedure: incidental learning phase

In the incidental learning phase, participants experienced high or low heat pain while being exposed to trial-unique object pictures (**Fig. 1a**; common to both the behavioral and fMRI experiments). In the fMRI study, this phase was conducted inside the fMRI scanner.

Importantly, the encoding of the object pictures was incidental (not instructed) in order to more closely resemble the incidental nature of encoding in many real-world situations. Regarding the incidental object pictures, participants were given slides with the following instructions in text: “In the middle of the screen you will see object pictures during the experiment that you can just look at.” Later, they were instructed: “Attention test: If the shapes or the object pictures blink, please press the spacebar. The object pictures are there to keep your attention on the screen during the heat stimulus.” Color pictures of objects were drawn from a database of images compiled via internet search (as used previously; Wimmer and Buchel, 2016); objects were largely composed of familiar non-food household items set on white backgrounds.

Heat pain was probabilistically cued (70% predictive) to allow for some prediction of pain but also for surprise at pain onset, with a design adapted from Atlas et al. (2010) (see also Geuter et al., 2017; Fazeli and Buchel, 2018). Across 4 blocks, 33 included high heat trials and 33 included low heat trials were presented (of 35 total; **Fig. 1a**), including 10 low-to-high and 10 high-to-low mismatch trials. To allow for initial adjustment to the task, data from two initial low heat trials were excluded. In the behavioral choice study, these objects were omitted from the choice phase or in the fMRI study, these objects were presented first and then excluded from analysis. Given the low number of mismatch trials and the relatively low and inconsistent effect of cues on ratings (see Results), all analyses focused on administered heat irrespective of cued heat to ensure reliability of imaging estimates.

To maintain attention on the screen during object presentation, participants were instructed to respond to occasional flickers in image brightness. The visual cue illumination flickered (decreased in illumination) once for 0.35 s in a random 50% of trials. Flicker timing was randomly distributed throughout the first 1.5 s of visual cue presentation. Similarly, in a separately determined random 50% of trials the object picture flickered in illumination during heat stimulation. When either a visual cue or object flicker was detected, participants were instructed to press the down button.

To detail the timing of events in an incidental learning phase trial, first, a visual cue signaling likely high or low heat was presented for 2.5 s. Participants responded to a visual flicker if one occurred. After a 4 s ISI, the incidental object appeared. The incidental object was presented for a total duration of 10 s. Participants responded to a visual flicker if one occurred. Following the heat stimulation and after a 4 s ISI, a pain rating scale appeared. Participants used left and right buttons to move a selection arrow from the initial cursor position (randomized between 4.5-5.5) to their experienced pain level and pressed the down button twice to make their selection; responses were self-paced. After the participant entered their response, trials were followed by a variable 2 s mean (range: 0.5-6 s) inter-trial-interval (ITI).

To allow for a better match between the appearance of the object and the onset of noticeable heat, heat onset started 0.75 s prior to object appearance (for a similar method, see Forkmann et al., 2013). The thermode temperature increased from baseline (33°C) to the desired temperature at a rate of 5 degrees per second, which translates to approximately 3.5 s to reach the range of the high heat temperature. After the 10 s object presentation period, the thermode temperature decreased at a similar rate. Thus, for low heat trials where the thermode did not need to reach a high value, the temperature during the 10 s presentation of the object was approximately constant at the desired value. For high heat trials, it took up to 2.5 seconds at the beginning of the 10 s period for the thermode to reach the peak. After each incidental learning phase block, the thermode was moved to a new location on the inner arm to avoid sensitization.

To maintain similar differences in subjective experience between the high and low heat conditions, temperatures were automatically adjusted throughout the task to maintain the targeted pain rating values. If the median of the previous 6 validly cued low heat trials fell below a rating of 1.5, the low temperature was increased by 0.2 °C; if the median rating was above 3, the low temperature was decreased by 0.2 °C. For the high temperature, if the median rating fell below 7.5, the high temperature was increased by 0.2 °C (if the temperature was below 50.5 °C). If a rating of “9” was given, indicating an intolerably high level of pain, the high temperature was decreased by 0.8 °C. Such on-line adjustments of administered temperature are not commonly employed in pain research that focuses on effects of expectation or placebo (e.g. Atlas et al., 2010), as in these cases administered temperature needs to be constant across the task. However, our focus here was on the subjective response to pain, and thus on-line adjustment allowed us to maintain very similar subjective responses to the majority of high and low heat stimuli.

Two pseudo-random orderings of incidental object pictures were used for counterbalancing object and heat associations. The assignment of abstract circles to high and low heat was also counterbalanced across participants. Further, after the first two blocks of the experiment, two new abstract circles were used as cues, with visual and verbal instruction about the new cues preceding the block. Visual cues were probabilistically associated with the level of heat, correctly predicting high or low heat in 70% of trials (Atlas et al., 2010). On invalid trials, the alternative heat level was administered. Additionally, 6 trials included a probe of cue-related pain expectancy: after 2.5 s of cue presentation, a question appeared below the cue asking participants whether they expected low or high heat to follow. These probes were used to ensure participants remained aware of the cue-pain associations. After the probe, trials continued as normal.

### Procedure: behavioral choice test phase

In the behavioral study, a surprise choice test phase followed the incidental learning phase to examine value memory for the objects incidentally associated with high or low heat in the preceding phase. Participants were instructed to select the object, out of two alternatives, that was associated with lower heat pain in the preceding phase. One object had been associated with the administration of high heat (independent of the cue) and the alternative object that had been associated with low heat (independent of the cue). Participants were instructed that they could win €0.50 euro for each correct choice of the lower heat object on top of their payment for participation.

The choices sampled each of the 66 objects from the incidental learning phase without repetition, resulting in 33 choices. The objects from the first two trials in the incidental learning phase were not included in any choice. Choices were presented in a pseudo-random order. A given choice included either 2 objects that had been correctly cued to be of low and high heat or a choice between one validly cued object and one invalidly cued object. We found no influence of the invalid cue or whether pain was higher or lower than expected on choice accuracy (p-values > 0.31) so we collapse across this factor in all analyses. Following these choices, an additional 4 trials presented choices between the abstract circle cues that had been predictive of high versus low heat pain.

To detail the timing of events on a choice trial, first, the choice options were presented serially in a random order (**Fig. 1b**). The first option was presented either on the left side or on the right side of the screen (determined at random) for 4 s, followed by a 1 s ISI. The second option was then presented in the alternate spatial location for 4 s, followed by a 1 s ISI. Then the first option returned to the screen, below the prompt “Lower heat? (€0.50 reward)”. Participants could select the on-screen option by pressing the ‘space’ key, or press the ‘left’ or ‘right’ key to alternate between the options. Alternation was allowed for an unlimited amount of time. After choice entry, a confidence rating followed, presenting the options: “Guess”, “Low”, “Medium”, and “High”. Participants responded using the 1-4 keys. A variable 3 s ITI followed.

### Procedure: fMRI memory test phase

In the fMRI study, a surprise memory test followed the incidental learning session. While collecting fMRI data, we assessed memory for the level of pain experienced with the object and recognition memory strength (**Fig. 1c**). Participants saw each of the “old” objects from the incidental learning phase. As noted above, the first two trials allowed for habituation and presented the first two objects from the incidental learning phase; these trials were not analyzed. The old objects were intermixed with 20 “new” objects for a total of 86 included trials.

The participants were given slides with text instructions, which included the following: “Try your best to indicate the heat strength that you remember being associated with them. It is likely that this is very difficult for you. Please just give your best guess or gut feeling. It is likely that you remember more than you think.” Before the start of this phase, they were reminded: “The heat question can seem difficult, but it’s very important to the experiment, so try to do your best. Guessing is okay!” For the recognition memory strength responses, participants were given the following instructions: “You have already seen most of the pictures in the first part, but some are also “new”. For the new pictures, it doesn’t matter what you say about the heat rating and the question of how sure you are.”

In our test phase trials, the pain association response was collected first, prior to the control measure of recognition strength, a reverse in question order compared to common in memory paradigms. This key feature was explicitly designed to allow us to best detect behavioral and neural evidence of pain value memory, and was motivated by multiple considerations. First, our design focused participants on any pain memory associations immediately upon the re-presentation of an object in order to increase behavioral performance and to best temporally isolate neural reactivation of pain, which we expected to be triggered immediately upon object re-presentation. Second, this order maximizes data collection by avoiding the loss of value memory responses for items rated as “new” as in common designs. Third, we prioritized value memory over the control recognition measure as our previous results found no link between reward value memory and recognition (Wimmer and Buchel, 2016). Fourth, our design facilitates generalization to decisions outside the lab: when making choices in the outside world, decisions about avoidance or approach can progress independently from recognition, and the ability to make such a determination quickly is likely to be adaptive.

To detail the timing of events on a memory phase trial, first, a single object was presented alone for 5 s. Next, after a 1 s ISI, an unmarked yellow-to-red heat scale with superimposed left- and right-pointing arrows was shown. Participants pressed the left or right buttons to indicate whether they thought that the object had been associated with low heat pain or high heat pain in the incidental learning phase. For objects that participants definitely considered to be “new”, participants were told that they could pick either the high or low heat response at random. If they were not sure if an object was new, participants were instructed to try to recall the level of heat it may have been paired with. All test phase responses were self-paced. Next, a confidence rating screen appeared with 4 levels of response: “guess”, “somewhat certain”, “certain”, and “very certain”. For stimuli participants believed were definitely new and thus had no associated heat experience, participants were instructed to respond with a low confidence answer. After a variable ISI (mean: 4 s; range: 3-6.5 s), a 6-point memory recognition strength scale was presented (e.g. Schwarze et al., 2012). Participants indicated whether they thought the object was “new” (not previously seen) or “old” (seen during the learning task) with 6 levels of response: “certain new”, “somewhat certain new”, “guess new”, “guess old”, “somewhat certain old”, “certain old”. Participants used the left and right buttons to move from the randomly initially highlighted “guess new” or “guess old” response option to their selected response and then pressed the down button twice to make their selection. A variable ITI with a mean of 4 s (range: 2-8 s) followed.

The order of the old pictures was pseudo-randomized from the incidental learning phase order, and the old and new pictures were pseudo-randomly intermixed. The duration and distribution of ITIs (or “null events”) was optimized for estimation of rapid event-related fMRI responses as calculated using Optseq software (http://surfer.nmr.mgh.harvard.edu/optseq/).

At the end of the experiment, participants completed a written questionnaire querying their knowledge of the task instructions and their expectations (if any) regarding the incidental object pictures. Task instructions and on-screen text were presented in German for all parts of the experiment; for the figures and methods, on-screen text has been translated into English.

### Data Acquisition

The experiment was presented using Matlab (Mathworks, Natick, MA) and the Psychophysics Toolbox (Brainard, 1997). For the behavioral study and the pain calibration phase of the fMRI study, data were collected using a 15” Apple Macbook Pro laptop. Responses were made using left and right arrow keys and the space key. In the scanner for the fMRI study, the task was projected onto a mirror above the participant’s eyes. Responses were made using a 4-button interface with a “diamond” arrangement of buttons. Skin conductance was recorded from the hypothenar of the left hand. The signal was amplified using a CED 2502 amplifier and digitized at 200 Hz using a CED micro1401 (both by Cambridge Electronic Design, Cambridge, UK) and downsampled offline to 100 Hz.

Whole-brain imaging was conducted on a Siemens Trio 3 Tesla system equipped with a 32-channel head coil (Siemens, Erlangen, Germany). Functional images were collected using a gradient echo T2*-weighted echoplanar (EPI) sequence with blood oxygenation level-dependent (BOLD) contrast (TR = 2460 ms, TE = 26 ms, flip angle = 80; GRAPPA factor of 2; 2 x 2 x 2 mm voxel size; 40 axial slices with a 1 mm gap). Slices were tilted approximately 30° relative to the AC–PC line to improve signal-to-noise ratio in the orbitofrontal cortex (Deichmann et al., 2003). Head padding was used to minimize head motion; no participant’s motion exceeded 3 mm in any direction from one volume acquisition to the next. For each functional scanning run, five discarded volumes were collected prior to the first trial to allow for magnetic field equilibration.

During the incidental learning phase, four functional runs of an average of 190 TRs (7 min and 48 s) were collected, each including 17 trials. During the memory test phase, four functional runs of an average of 196 TRs (8 min and 2 s) were collected, each including 22 trials. If a structural scan had been collected for the participant at the center within the past 6 months, the previous structural scan was used. If not, structural images were collected using a high-resolution T1-weighted magnetization prepared rapid acquisition gradient echo (MPRAGE) pulse sequence (1 x 1 x 1 mm voxel size) between the incidental learning phase and the immediate memory test phase (12 participants). We found no relationship between the consequently varying time delay between the end of the incidental learning phase and the start of the test phase on value memory performance, recognition memory performance, or fMRI pain reactivation measures (from n = 28 participants with timing and structural scan origin information).

All voxel locations are reported in MNI coordinates, and results are displayed overlaid on the average of all participants’ normalized high-resolution structural images using the xjView toolbox (http://www.alivelearn.net/xjview) or AFNI (Cox, 1996).

### Behavioral Analysis

Our primary behavioral question was whether memory-based decisions were influenced by the pain that had been experienced with objects in the preceding incidental learning phase. In the behavioral experiment, choice trials were excluded if the administered heat for the high heat stimulus did not exceed that for the low heat stimulus (in rare cases when the thermode failed to increase temperature; on average less than 1 trial per participant).

In both experiments, we conducted simple a priori comparisons of behavioral performance to chance (50%) using t-tests, with a significance threshold of p < 0.05 (two-tailed). We also examined the influence of cue expectation on pain ratings using a paired t-test. In the fMRI experiment, we further verified in initial comparisons that “old” objects (whether paired with high or low pain) were recognized at a higher rate than “new” objects.

To further investigate value memory, multilevel regression models were implemented in R using lme (from the nlme package) for linear regression and glmmTMB (from the glmmTMB package) for logistic regression. All predictors and interactions were included as random effects, following the ‘maximal’ approach (Barr et al., 2013). In all regressions, participant was entered as a random effect along with all other variables of interest. Correlations between random effects were included when convergence was achievable with this structure. All reported p-values are two-tailed. In a control model, we verified that the presence vs. absence of a visual “flicker” during object presentation was not related to value memory or recognition memory strength.

We additionally tested whether nonsignificant results were weaker than a moderate effect size using the two-one-sided t-test (TOST) procedure (Schuirmann, 1987; Lakens, 2017) and the TOSTER library in R (Lakens, 2017). In the behavioral experiment (n = 21), we used bounds of Cohen’s *d* = 0.64, where power to detect such a medium-size is estimated to be 80%. In the larger fMRI sample (n = 29), we used bounds of Cohen’s *d* = 0.55 to achieve the same estimated power.

### fMRI preprocessing

Preprocessing and data analysis were performed using Statistical Parametric Mapping software (SPM12; Wellcome Department of Imaging Neuroscience, Institute of Neurology, London, UK). Before preprocessing, slices with artifacts were identified as having mean image intensity greater than or equal to 5% above the across-run slice mean. Individual slices with artifacts were replaced with the mean of the two surrounding timepoints using a script adapted from the ArtRepair toolbox (Mazaika et al., 2009). Images were then slice-timing corrected, realigned to correct for participant motion, and then spatially normalized to the Montreal Neurological Institute (MNI) coordinate space by estimating a warping to template space from each participant’s anatomical image and applying the resulting transformation to the EPIs. Images were filtered with a 128 s high-pass filter and resampled to 2 mm cubic voxels. Images were then smoothed with a 6 mm FWHM Gaussian kernel for univariate and connectivity analyses.

### fMRI univariate analyses

fMRI model regressors were convolved with the canonical hemodynamic response function and entered into a general linear model (GLM) of each participant’s fMRI data. The six scan-to-scan motion parameters produced during realignment were included as additional regressors in the GLM to account for residual effects of participant movement. All regressions were conducted with automatic orthogonalization in SPM turned off.

Our primary univariate analysis was a “localizer” analysis to identify main effects of pain during the incidental learning phase. The GLM included regressors for the cue period (2.5 s duration), the initial pain onset period (2 s), the full pain and object presentation period (10 s), and the pain rating period (with a variable duration based on response time). The cue period regressor was accompanied by a parametric modulator contrasting high versus low expected pain. The pain onset period regressor was accompanied by two parametric modulators: the mismatch between cue and pain as well as the unsigned (absolute value) mismatch between cue and pain (these regressors were not correlated; r = 0.007). The full pain period regressor was accompanied by a parametric modulator representing the pain rating given on that trial. Note that the regions identified as correlating with pain during the 10 s pain period were the same with or without the inclusion of the 2 s pain onset regressor.

Then we conducted several control univariate analyses. First, we examined learning phase activity correlated with later successful pain value memory and recognition memory strength. This model was based on the GLM above, but instead of the pain rating parametric modulator, we included parametric modulators for subsequent correct value memory and subsequent recognition memory strength. Separate parametric regressors were used for high and low pain-associated objects to allow for baseline differences (yielding four parametric regressors in total); results were then combined at the second level.

The remaining control univariate analyses examined activity in the test phase. In these models, a 5 s regressor modeled activity during the object re-presentation period. Additional regressors modeled the pain memory response period, the pain memory confidence response period, and the memory response period; the durations for all these periods matched the participant’s response time. First, we looked for univariate correlates of pain reactivation to confirm that any multivariate results were not primarily driven by univariate activity. Thus, the object re-presentation regressor was accompanied by a parametric modulator representing the level of heat pain experienced with objects in the preceding learning phase. Second, a control test phase univariate analysis examined correlates of pain value memory success and recognition memory strength. Here, the object re-presentation regressor was accompanied by parametric regressors representing value memory success and recognition memory strength; as in the learning phase, separate regressors were used for high and low pain-associated objects and were combined at the second level.

### Multivariate fMRI analyses

To test our primary fMRI prediction that patterns of BOLD activity associated with negative emotional experience were reactivated at retrieval, we utilized multivariate classification analyses. These analyses used the non-smoothed fMRI data. In the incidental learning phase and the memory test phase we estimated mass-univariate GLMs where each trial was modeled with a separate regressor. For the incidental learning phase, each regressor modeled the onset of an object and continued through the 10 s duration of the heat stimulus. For the memory test phase, each regressor began at the onset of the object and continued for the 5 s duration of object presentation (prior to any responses). Models included the six nuisance motion regressors (translations and rotations).

Multivariate analyses were conducted using The Decoding Toolbox (Hebart et al., 2014). Classification utilized a L2-norm learning support vector machine (LIBSVM; Chang and Lin, 2011) with a fixed cost of *c* = 1. The classifier was trained on the full incidental learning phase balanced via bootstrapping. The trained classifier was then tested on the full memory test phase data. Note that for the primary across-phase classification analysis, no cross-validation is necessary for training because no inferences are drawn and no results are reported from the incidental learning phase data. Memory test phase classification is reported as area under the curve (AUC), which uses graded decision values and better accounts for biases in classification that may arise due to the different processes engaged by the incidental learning and memory test phases. Supplemental ROI analyses examined training and testing within the learning phase or memory test phase using cross-validation. Using cross-validation, we computed the strength of discriminability in the localizer phase in our regions of interest.

Additionally, we conducted a searchlight analysis for further localization using The Decoding Toolbox (Hebart et al., 2014). We used a 4-voxel radius spherical searchlight (approx. 208 voxels). Training of the classifier on the incidental learning phase and testing on the memory test phase were conducted as described above for the ROI MVPA analyses. Individual subject classification accuracy maps were smoothed with a 6 mm FWHM kernel before group-level analysis. We also performed covariate analyses to determine whether behavioral performance was correlated with classification accuracy.

It has been shown that it is not valid to conduct statistical inference specifically on cross-validated classification accuracy measures of information using *t*-tests (Allefeld et al., 2016). In part, as informational measures cannot be below zero, assumptions underlying the *t*-test are violated for cross-validation within the same dataset. Our classifier training and testing were conducted on separate datasets (“cross-classification” between the incidental learning and the memory test phase) which does allow for potential “true” below-zero values, a case not addressed by Allefeld et al. (2016). Further, we found that cross-classification AUC values in all our regions of interest followed a normal distribution (Anderson-Darling goodness-of-fit hypothesis test). While the above concern may still apply to inferences made about the main effects of pain during the incidental learning phase, our primary hypothesis rests on the cross-classification of pain-related patterns from the memory test phase.

### Connectivity analyses

We additionally conducted psychophysiological interaction (PPI) analyses to examine differences in functional connectivity for successful versus unsuccessful value memory retrieval. These analyses used a hippocampal ROI as the seed region (defined in Results). In the incidental learning phase, we estimated a PPI contrasting correct versus incorrect later value memory retrieval, modeling the 10 s duration of the object and pain period. In the memory test phase, we estimated a similar PPI analysis, contrasting correct versus incorrect value memory retrieval, modeling the 5 s duration of the object presentation period. At the second level, we performed correlation analyses to determine whether behavioral performance was related to differences in connectivity for correct versus incorrect encoding or retrieval of value memory associations.

### Statistical correction and regions of interest

For both univariate and searchlight results, linear contrasts of univariate SPMs were taken to a group-level (random-effects) analysis. We report results corrected for family-wise error (FWE) due to multiple comparisons (Friston et al., 1993). We conduct this correction at the peak level within small volume ROIs for which we had an a priori hypothesis or at the whole-brain cluster level (in each case using a cluster-forming threshold of p < 0.005 uncorrected, except for the pain rating correlation, where we used p < 0.00001 to yield more interpretable clusters).

We focused on two a priori ROIs motivated by two separate hypotheses. Given the anterior insula’s role in processing the affective qualities of pain (Kurth et al., 2010; Wiech et al., 2014), we predicted that the insula may relate to the modulation of memory by pain. For this pain hypothesis-motivated anterior insula ROI, we first created a bilateral anterior insula mask (Brooks et al., 2002; Wiech et al., 2014), covering the insular cortex anterior to y = 9, as well as up to 4 millimeters lateral or superior to the insular cortex to account for signal blurring and anatomical variability. This mask was further restricted by the main effect of pain rating taken from the incidental learning phase localizer GLM defined above, thresholded at p < 0.0001 uncorrected (https://neurovault.org/collections/6126/). We also defined a broader pain-related mask based on the localizer GLM thresholded at p < 0.0001 uncorrected, excluding the cerebellum. Separately, we focused on the hippocampus because of its role in episodic and relational memory (Eichenbaum and Cohen, 2001; Davachi, 2006). We also conducted follow-up analyses in the anterior hippocampus, given its role in negative emotion-related memory and generalization (Fanselow and Dong, 2010; Poppenk et al., 2013). The bilateral hippocampus ROI was derived from the Harvard-Oxford atlas at a threshold of 50%. We focused on a restricted mask of the hippocampus in order to limit the size of the ROI for multivariate analyses. We confirmed that there was no overlap between the hippocampus and pain-related masks. We defined the anterior hippocampus as the mask region anterior to Y = −21, approximating the position of the uncal apex (Poppenk et al., 2013). While somatic processing of thermal pain does not primarily involve the amygdala, as a control we also examined the amygdala, defined from the Harvard-Oxford atlas at a threshold of 50%.

Correlations between classification accuracy and behavioral performance were conducted using Pearson’ correlation. Statistical comparison of the difference between correlations was computed using Steiger’s test for differences in dependent correlations.

### Data availability

Behavioral data are available on the Open Science Framework (https://osf.io/gr9xd/). Whole-brain fMRI results are available on NeuroVault (https://neurovault.org/collections/6126/).

## Results

### Choice study behavior

Our behavioral question was whether single aversive episodes can lead to memory for value associations, which we refer to as value memory, and that such memory supports later decision making. In the behavioral study, participants experienced episodes of low or high heat pain incidentally associated with trial-unique object pictures. Subsequently, in a surprise choice test phase, participants made choices between two objects that had been incidentally associated with different levels of heat, with the goal of choosing the object that had been paired with low heat.

During the incidental learning phase, participants could clearly discriminate the heat pain levels: on the 1-8 rating scale, where ‘8’ corresponds to high pain, the mean pain rating for high pain stimuli was 7.00 (95% CI [6.68 7.31]), while the mean pain rating for low pain stimuli was 2.26 [1.94 2.57]. Participants’ pain ratings were also highly correlated with the administered heat temperature on a trial-to-trial basis (mixed-effects model coefficient β = 0.9399 [0.7682 1.036]; z = 10.899 p < 0.001). The cue preceding the high or low pain was inaccurate on 30% of trials. We found no significant interaction between high versus low pain and cue validity (β = 0.0292 [−0.0295 0.0867]; t = 1.010, p = 0.326). There was no significant effect of invalid cues on low pain ratings (valid 2.23 [1.94 2.52]; invalid 2.32 [1.93 2.72]; β = 0.0472 [−0.0349 0.1304]; t = 1.106, p = 0.282) and a no significant effect of invalid cues on high pain ratings (valid 7.00 [6.69 7.32]; invalid 6.98 [6.64 7.33]; β = −0.0111 [−0.0893 0.0655]; t = −0.287, p = 0.821). The minimal influence of the cue on ratings is likely due to the use of two very different and easily discriminable temperatures, which differs from previous work (Atlas et al., 2010; Fazeli and Buchel, 2018).

In the incentivized choice test phase, participants were successfully able to choose the low pain object over the high pain object (mean 58.7% correct choices [53.2 64.2]; versus chance (50%), t_(20)_ = 3.28, p = 0.0037). Interestingly, we found that choice performance significantly increased with the difference in the learning phase pain ratings between the two choice objects (β = 0.1278 [0.0228 0.2327]; z = 2.387, p = 0.0170; **Fig. 2a**). Choice performance also increased with higher levels of choice confidence (β = 0.4409 [0.2506 0.6312]; z = 4.541, p = 0.000006), indicating significant metacognitive awareness. In this experiment, a supplemental recognition strength measure was not collected. Thus, we could not examine any links between choices and recognition, which may be related to attention during incidental encoding. We would expect inattention during learning to decrease performance, as participants would need to rely on information encoded about the alternative object. However, we speculate that instances of inattention for one or both objects would be reflected in low confidence, low accuracy choices, which would, if anything, decrease our ability to detect an effect. Overall, the results from the behavioral study demonstrate that value-based choices, here to avoid a pain-associated item, can be guided by the strength of single experiences.

**Figure 2.**
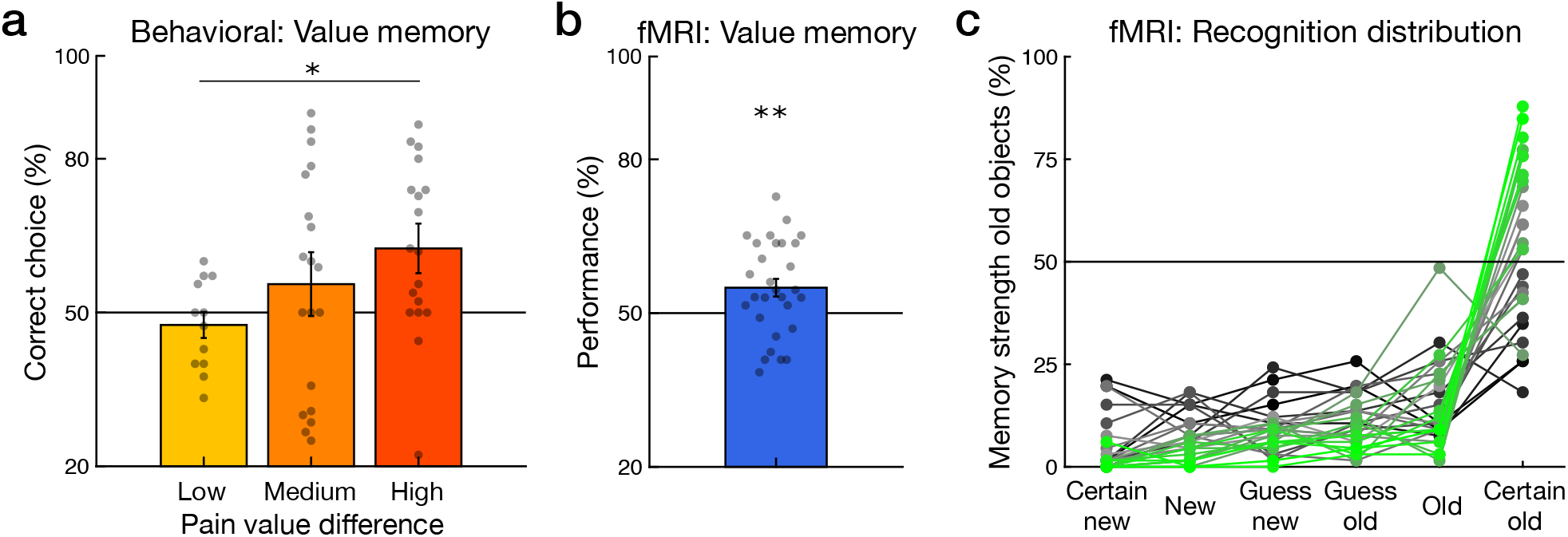
Decision making and value memory performance. ***a***, In the behavioral experiment, accuracy in selecting the object that had been incidentally associated with low versus high pain was significantly related to the difference in pain reported for the objects during the incidental learning phase (regression on continuous measure). For visualization only, the pain rating difference between choice options was binned based on whether the options differed by <= 3 rating points (Low), > 3 and <= 5 points (Medium), and 5 or more points (High). ***b***, Test phase performance in the fMRI study. Participants exhibited value memory: memory for the value associated with single episodes. ***c***, Recognition memory response distribution for old objects for individual participants (n = 28 with ratings data). Participants are colored based on memory performance; green for higher rates of combined “old” and “certain old” responding and black for lower rates. (Individual points represent individual participants; error bars represent standard error of the mean (SEM); * p < 0.05; **p < 0.01.)

### fMRI study behavior

In the fMRI study, as in the behavioral study, our behavioral question was whether single aversive episodes can support later value-based decision making. Participants experienced episodes of high or low heat pain incidentally associated with trial-unique object pictures. Subsequently, in a surprise memory test, participants were cued with an object and instructed to remember whether the object was associated with high or low pain in the preceding incidental learning phase. Following this key pain value memory response, as a control, participants then rated their recognition strength for the object.

In the incidental learning phase, pain ratings given after each trial reliably differentiated high and low heat (high, 7.34 95% CI [7.20 7.48]; low, 2.34 [2.13 2.55]; scale range: 1-8). Participants’ pain ratings were highly correlated with the administered heat temperature on a trial-to-trial basis (β = 0.7093 [0.6406 0.7799]; z = 20.12, p < 0.001). The cue preceding the high or low pain was inaccurate in 30% of trials. We found a significant interaction between high versus low pain and cue validity (β = 0.0870 [0.0428 0.1310]; t = 3.967, p < 0.001), unlike the behavioral study. This interaction was driven by a positive effect of invalid cues on low pain ratings (valid 2.27 [2.09 2.44]; invalid 2.52 [2.19 2.85]; β = 0.1260 [0.0287 0.2264]; t = 2.461, p = 0.008) and a numerically negative effect of invalid cues on high pain ratings (valid 7.37 [7.24 7.50]; invalid 7.27 [7.10 7.45]; β = −0.0480 [−0.1047 0.007]; t = −1.686, p = 0.094).

In the surprise memory test, we found that value memory accuracy was significantly above chance (54.96% correct [51.43 58.49]; t_(28)_ = 2.879, p = 0.0076; **Fig. 2b**). Importantly, value memory accuracy significantly increased with increasing confidence (β = 0.2979 [0.1689 0.4270]; z = 4.525, p = 0.000006; n = 28 participants with confidence and memory ratings), with performance rising to 72.42% at the highest confidence level. As in the behavioral study, this relationship indicates that value memory responses were often based on underlying accurate memories that were accessible to awareness. Nevertheless, the level of performance was lower than what we observed in a study where objects were incidentally associated with monetary reward (61%; Wimmer and Buchel, 2016). We did not find better value memory performance for objects associated with more extreme high or low pain ratings (high pain rating difference between correct versus incorrect episodes, p = 0.341; low pain rating difference, p = 0.753), unlike the behavioral choice experiment. It is possible that the binary choice measure in the behavioral study was more sensitive to this effect.

We then examined the control measure of recognition memory strength. Recognition responses were collected after the pain value memory confidence responses on each trial. Participants reliably discriminated old from new objects (old object mean 4.87 [4.65 5.09]; new object mean 2.11 [1.85 2.37], p < 0.001), with a recognition rate of 79.90% and relatively high corrected rate of 66.70% (hits minus false alarms; range 19.48 – 98.51) (Wimmer and Buchel, 2015). More than two-thirds of old items were rated as ‘old’ or ‘certain old’ (69.0% [63.0 75.1]). As illustrated in the response distribution for all participants in **Fig. 2c**, recognition memory for objects experienced in the learning phase were highly biased toward ‘old’ responses (percent of the 66 old trials rated ‘certain new’ = 4.3%; ‘new’ = 7.1%; ‘guess new’ = 8.9%; ‘guess old’ = 10.7%; ‘old’ = 15.0%; ‘certain old’= 54.0%).

We found that recognition memory strength was not modulated by heat pain experienced in the preceding phase (where a rating of ‘6’ represents certain old: high pain 4.86 [4.65 5.07]; low pain 4.87 [4.65 5.09]; two-one-sided equivalence test (TOST) p = 0.006; thus we can reject the presence of a medium- or larger-sized effect; n = 28 participants with value memory confidence and recognition memory ratings). The null effect of pain on recognition memory was validated in a multilevel regression model (β = 0.0096 [−0.051 0.070]; z = 0.311, p = 0.756; TOST p = 0.007). Recognition memory strength was also not significantly related to value memory accuracy (β = 0.0542 [−0.0165 0.1250]; z = 1.501, p = 0.133), though we cannot rule out a medium-sized effect (TOST = 0.078). In a combined model including trial-by-trial value memory confidence alongside recognition, value memory confidence remained significantly related to value memory accuracy (p = 0.000008), while recognition was not significantly related to value memory accuracy (p = 0.191; TOST p = 0.055). Thus, we find no relationship between experienced heat and recognition memory, and no relationship between recognition memory and value memory performance.

The results from both the behavioral and fMRI experiments demonstrate that single aversive episodes of high or low heat pain can support later memory-based decisions. When making a choice between two aversive options that have been only experienced once before (Kahneman et al., 1993), a value-based decision is, from a different perspective, a decision about remembered stimulus intensity. Critically, however, in the domain of aversive experiences, which carry a negative valence, a decision about stimulus intensity is inherently a decision about value.

Importantly, arousal (or stimulus intensity) alone is unlikely to primarily drive these behavioral results. In a previous study, we combined the same incidental learning phase procedure from the current study with a reward-based incidental learning phase (Wimmer and Buchel, 2016). At a surprise test, participants chose between reward-versus pain-associated objects or between two objects within the same valence. If value memory choice performance was based primarily on arousal, then performance would be poor when choosing between high-arousal reward-associated objects and high-arousal pain-associated objects. However, in new analyses, we found that performance on choices between high reward- and high pain-associated objects was, if anything, higher than performance on choices between two objects with the same valence (reward versus pain 69.09% [63.67 74.51]; t_(19)_ = 7.376, p < 0.0001; same-valence (within reward or pain) 62.50% [54.55 70.45]; t_(19)_ = 3.290, p = 0.004; difference t_(19)_ = 1.972, p = 0.063).

### fMRI univariate pain results

In the imaging analyses, we first examined whether heat pain activated the network of regions implicated in pain processing (Apkarian et al., 2005; Tracey and Mantyh, 2007). We found that trial-by-trial pain ratings positively correlated with activation in regions previously associated with pain processing including the anterior and posterior insula, cingulate, thalamus, and secondary somatosensory cortex (all p < 0.05 whole-brain FWE corrected; **Fig. 3**). A region of the right hippocampus also showed a correlation with pain ratings (20, −18, −14; z = 3.55, p = 0.025 SVC), although this effect could be related to the use of spatial smoothing in the data underlying the univariate analyses; note that subsequent multivariate analyses use non-smoothed data. We also examined the response to pain-predictive cues. We found activation for high versus low cues in a cluster extending from the left anterior orbitofrontal cortex (OFC) to more posterior medial OFC (−32, 50, −14; z = 4.12; p < 0.001 whole-brain FWE; no other regions survived whole-brain correction; https://neurovault.org/collections/6126/), but no significant activation in pain-related regions or the hippocampus. No regions exhibited significantly greater activity for low versus high pain cues.

**Figure 3.**
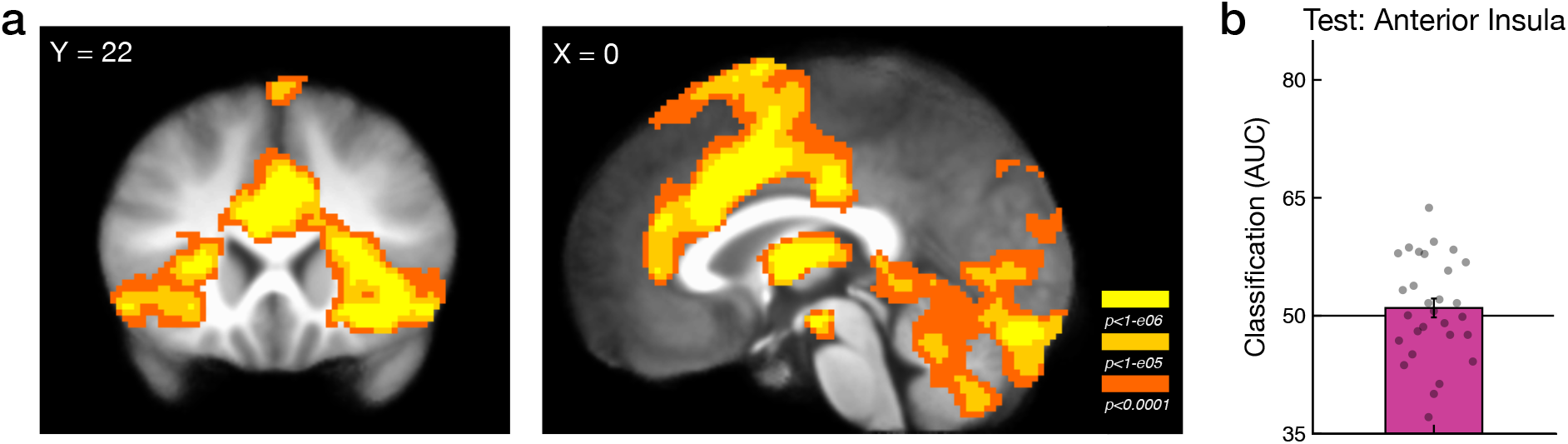
Heat pain response during the incidental learning phase, and heat pain reactivation in the insula during re-presentation of pain-associated objects. ***a***, Pain rating correlation during heat pain administration in the anterior insula and other regions (images thresholded at p < 0.0001, activation significant at p < 0.05 FWE; unthresholded map available at https://neurovault.org/images/306227/). ***b***, Classification of later re-presentation of high- versus low-pain objects in the memory test phase based on patterns of activation to pain in the anterior insula pain-responsive region of interest. (Individual points represent individual participants; error bars represent SEM.)

### fMRI multivariate results

Next, we addressed our primary question of whether distributed patterns of activity during object re-presentation reflected incidental pain value associations. We trained a classifier on the multivoxel patterns of activation evoked by actual pain in the incidental learning phase. To check whether the classifier trained on actual pain experience during the incidental learning phase was able to classify pain, we examined cross-validated results in regions of interest defined by the univariate correlation with pain ratings. We defined two masks, one including voxels in the anatomical anterior insula that exhibited a correlation with pain ratings (using an uncorrected p < 0.0001 threshold) and one including any brain voxels correlated with pain ratings (p < 0.0001, uncorrected). MVPA analyses revealed high rates of classification of high versus low pain in the anterior insula (84.5% AUC classification performance; note that these results are provided for illustration only given that the definition of the ROI was itself based on pain responses) and from the whole-brain pain region mask (89.1%). We also found that distributed activity patterns in the hippocampus discriminated high versus low pain (68.4%, p < 0.0001). Note that the data underlying the multivariate analyses are not spatially smoothed, making it unlikely that effects in adjacent regions contribute to multivariate results.

Building on the behavioral finding that single aversive experiences can support memory-based decisions, we then turned to our primary question of whether patterns of neural activity during high versus low heat pain exposure were reactivated when participants were re-presented with heat-paired object pictures. During the memory retrieval phase, participants were presented with an object for 5 s, followed by a heat rating prompt where they responded with whether they remembered that the object picture had been paired with high versus low heat (**Fig. 1c**). Using the multivoxel p classifier trained on the activation evoked by actual pain in the incidental learning phase, we then tested the performance of this classifier on activation during object representation.

Upon re-exposure to objects incidentally paired with heat pain, we found no significant evidence for reactivation of pain-related patterns in traditional pain-processing regions, including the anterior insula (**Fig. 3a**). Classification performance in the anterior insula was not greater than chance (51.43 AUC [48.84 53.51]; t_(28)_ = 1.07, p = 0.312; TOST p = 0.032; **Fig. 3b**). Further, in a network of regions across the whole brain that exhibited a correlation with pain experience, classification performance at test was also not greater than chance (51.43 [48.70 54.17]; t_(28)_ = 1.07, p = 0.293; TOST p = 0.035). We predicted that somatic sensation (heat) would primarily be reflected in the insula, but we also examined activity in the amygdala as a control region. Amygdala patterns of activity did not show evidence of reactivation of pain associations (50.61 [47.52 53.71]; t_(28)_ = 0.41, p = 0.688; TOST p = 0.008).

In the hippocampus, however, we found evidence for significant reactivation of pain-related patterns (53.31 [50.30 56.32]; t_(28)_ = 2.25, p = 0.032; **Fig. 4a**). As noted above, value memory behavioral performance in the current experiment was relatively low; thus, we also examined a subgroup of participants that approximated the stronger behavioral value memory performance in our previous study using reward (Wimmer and Buchel, 2016). Within a subgroup of 21 participants who exhibited value memory performance above 50% (mean 59.5% performance), we found numerically stronger classification of pain-associated episodes in the hippocampus (55.07 [51.38 58.77]; t_(20)_ = 2.87, p = 0.0096). This brain classification performance is of the same magnitude as previously reported for reward episode classification (Wimmer and Buchel, 2016).

**Figure 4.**
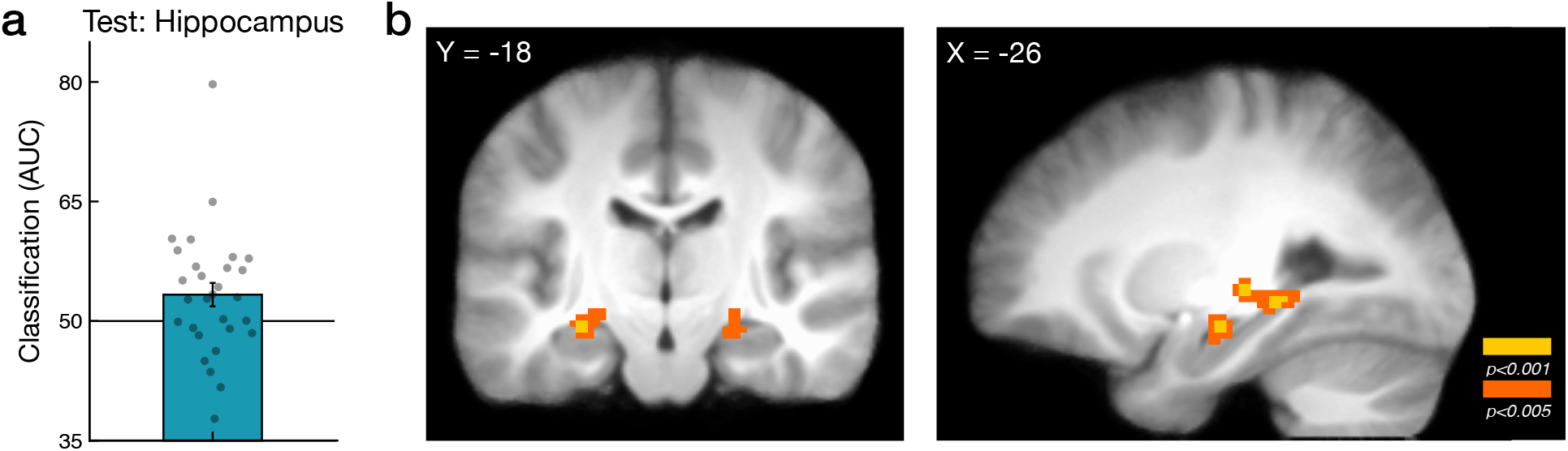
Heat pain reactivation in the hippocampus during re-presentation of objects. ***a***, Significant classification of later re-presentation of high versus low pain objects in the test phase based on incidental learning phase patterns of activation to pain in the hippocampus (p = 0.032). (Individual points represent individual participants; error bars represent SEM.) ***b***, Illustration of positive but non-significant searchlight reactivation of high versus low pain in the hippocampus. Images thresholded at p < 0.005 uncorrected for display, clusters p < 0.10 SVC. (For unthresholded map, see: https://neurovault.org/images/390597/.)

We tested for but did not find a difference in classification accuracy in the hippocampus based on whether participants were correct in their pain memory response (correct high pain vs low: 54.56 [49.93 59.20]; *t*_(28)_ = 2.018, p = 0.053; incorrect high pain vs low: 52.31 [48.32 56.31]; *t*_(28)_ = 1.186, p = 0.26; comparison, *t*_(28)_ = 0.75, p = 0.46). If such a difference in classification due to correct behavioral responses had been found, it would have been difficult to distinguish actual value memory reactivation from an effect of behavioral response (high versus low pain) in the test phase that itself triggered an affective reaction. This null effect of accuracy replicates a previous null result for reward associations in reward-responsive regions (Wimmer and Buchel, 2016). We also verified that the reactivation effect was not driven by a simple effect of test phase pain memory response itself. A classifier trained on pain and tested on test phase pain memory response (high vs low) found no effect (50.20 [46.95 53.44]; *t*_(28)_ = 0.13, p = 0.90; TOST p = 0.004).

We next confirmed that the pain pattern reactivation result in the hippocampus was selective to distributed multivariate patterns and not overall changes in activity. We extracted test phase trial-by-trial univariate beta values, averaged across the hippocampus, and trial-by-trial classifier decision values, representing the strength of evidence for high versus low heat pain reactivation in the hippocampus. First, in a multilevel regression analysis utilizing trial-by-trial classifier decision values from the multivariate analysis, we validated the finding that test phase high versus low pain reactivation in the hippocampus was significantly predicted by high versus low pain experience in the incidental learning phase (β = 0.0415 [0.0104 0.0727]; t = 2.613, p = 0.009). Interestingly, we found a numerically stronger link between subjective pain ratings and pain reactivation (β = 0.0182 [0.0064 0.0300]; t = 3.037, p = 0.0024). Second, in control analyses we found that test phase univariate activity at object representation was not related to pain experienced in the learning phase (β = 0.0084 [−0.0302 0.04693]; t = 0.427, p = 0.669; TOST p = 0.009). In a combined model, we found that multivariate pain patterns were significantly related to high versus low experienced pain while univariate activation was not (multivariate β = 0.1622 [0.0357 0.2886]; z = 2.514, p = 0.0119; univariate β = 0.0305 [−0.1045 0.1656]; z = 0.443, p = 0.658; TOST p = 0.009). Further, the trial-by-trial multivariate hippocampal reactivation measure itself was unrelated to trial-by-trial hippocampal univariate activity (p = 0.702).

To demonstrate the selectivity of pain pattern reactivation to episodic high versus low heat pain experiences, we tested for a relationship between pain pattern reactivation and test phase recognition memory strength. As value associations can drive behavior independent of explicit memory (e.g. Wimmer and Shohamy, 2012), and following a previous null finding relating recognition and reactivation of episodic reward associations (Wimmer and Buchel, 2016), we did not expect pain pattern reactivation to be related to memory strength. First, we found that classifier decision values indexing pain reactivation were not significantly related to graded recognition memory strength ratings (β = 0.0193 [−0.0041 0.0428]; t = 1.617, p = 0.106; TOST p = 0.095; n = 28). Next, in a model including both high versus low pain and recognition memory as independent variables, we found that high versus low pain remained significantly related to pain reactivation in the hippocampus (p = 0.0219) while the relationship between recognition memory and reactivation remained non-significant (p = 0.110). We also found no significant relationship between pain reactivation and the interaction between high versus low pain and recognition memory (β = 0.0192 [−0.0025 0.0411]; t = 1.733, p = 0.083). Finally, we found no relationship between reactivation memory and the absolute value of multivariate pain pattern reactivation (a measure of the strength of classifier evidence in either direction; β = −0.0182 [−0.0389 0.0024]; t = −1.733, p = 0.083). Together, these control analyses support the interpretation that our results are selectively related to pain value memory and not the strength of recognition memory for an experience.

The above classification analyses demonstrated that distributed patterns of activity in the hippocampus but not pain-related regions showed significant classification of pain reactivation. To examine classification performance based on local information, we performed a searchlight analysis (Kriegeskorte et al., 2006). This analysis revealed no significant clusters across the whole brain, no effects in the insula or wider pain-related ROI mask, and no significant effects in the hippocampus. However, illustrating local information that may drive the whole-ROI result above, three clusters in the hippocampus showed non-significant positive effects (left posterior: −26, −36, −4; z = 3.35, p = 0.089 SVC; right middle: 24, −24, −12; z = 3.34, p = 0.091 SVC; left anterior: −26, −16, −14; z = 3.71, p = 0.097; **Fig. 4b**; unthresholded map available at https://neurovault.org/collections/6126/).

### Multivariate reactivation and value memory performance

We then examined the critical question of whether individual differences in pain-related reactivation was related to participants’ pain value memory performance. We correlated the whole-brain searchlight analysis results with individual performance in value memory retrieval. A region in the left anterior hippocampus showed a significant relationship between searchlight pain classification strength and value memory performance (−28, −12, −26; z = 3.64, p = 0.038; **Fig. 5a**). This correlation was also evident in an ROI analysis of the anterior hippocampus (r = 0.470, p = 0.0204, corrected for two comparisons; **Fig. 5b**). Comparisons of the anterior and posterior hippocampus ROIs showed a stronger correlation with behavior in the anterior versus posterior hippocampus (posterior r = −0.108, p > 1.0, corrected for two comparisons; difference z = 2.36, p = 0.018).

**Figure 5.**
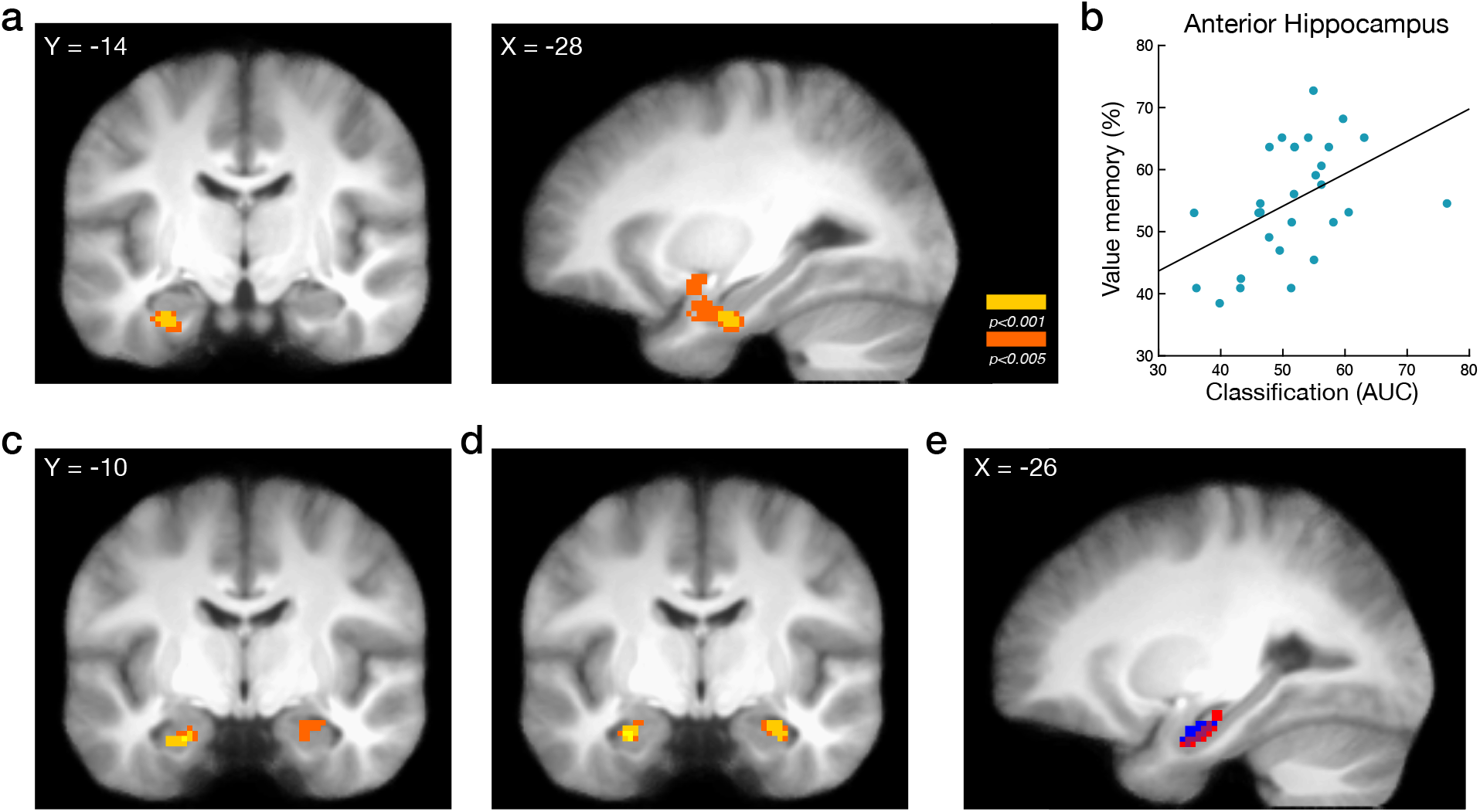
Hippocampal classification and the relationship between hippocampal connectivity and behavior. ***a***, In a searchlight analysis, the reactivation of pain patterns in the left anterior hippocampus was positively associated with value memory behavioral performance across participants (full anterior MTL cluster selected for display; images thresholded at p < 0.005 for display, p = 0.038 SVC; https://neurovault.org/images/390598/) ***b***, Illustration of the anterior hippocampus reactivation-performance relationship in an anatomical anterior hippocampal mask. Individual points represent individual participants. ***c***, Connectivity between the anterior hippocampus for correct versus incorrect value memory retrieval trials correlated with value memory performance during the incidental learning phase (left hippocampus p = 0.022 SVC; right hippocampus p = 0.084 SVC; https://neurovault.org/images/390601/), and ***d***, during the memory test phase (left hippocampus p = 0.003 SVC; right hippocampus p = 0.035 SVC; https://neurovault.org/images/390602/). ***e***, Conjunction in the left hippocampus of the connectivity relationship with individual differences in value memory performance in the incidental learning phase (red), memory test phase (blue) and overlap (purple).

The brain-behavior correlation in the anterior hippocampus was selective to the multivariate pain reactivation measure and to value memory performance. First, differential univariate anterior hippocampal activity for re-exposure to high versus low pain objects was not related to value memory performance (p = 0.353). Second, we found no link between learning phase pain discrimination in the anterior hippocampus and test phase value memory performance (r = −0.239, p = 0.213). Importantly, variability in test phase pain pattern reactivation in the anterior hippocampus was also unrelated to recognition memory (r = 0.298, p = 0.132). Further, even though value memory and recognition memory measures were positively correlated across participants, in a model including both measures as independent variables, value memory performance remained significantly related to anterior hippocampus pain reactivation (t_(25)_ = 2.249, p = 0.034) while there was no relationship with recognition memory (t_(25)_ = −0.358, p = 0.723).

These results demonstrate that for single aversive episodes, distributed multivariate patterns in the hippocampus during object re-presentation significantly resemble those evoked by actual pain during the original experience. More generally, this demonstrates that affect-related neural patterns are re-expressed at later retrieval.

### Connectivity and value memory performance

We then examined whether connectivity during incidental encoding or at test was related to value memory. Specifically, we tested for differences in connectivity during successful versus unsuccessful value memory using a PPI analyses, focusing on the hippocampus. Prior to the PPI analyses, we examined univariate correlates of successful pain value memory. During the incidental learning phase, we found no significant positive correlations with value memory accuracy (**Table 4-1**). In the test phase, we also found no activity significantly correlated with value memory accuracy (**Table 4-2**).

**Table 3-1.**
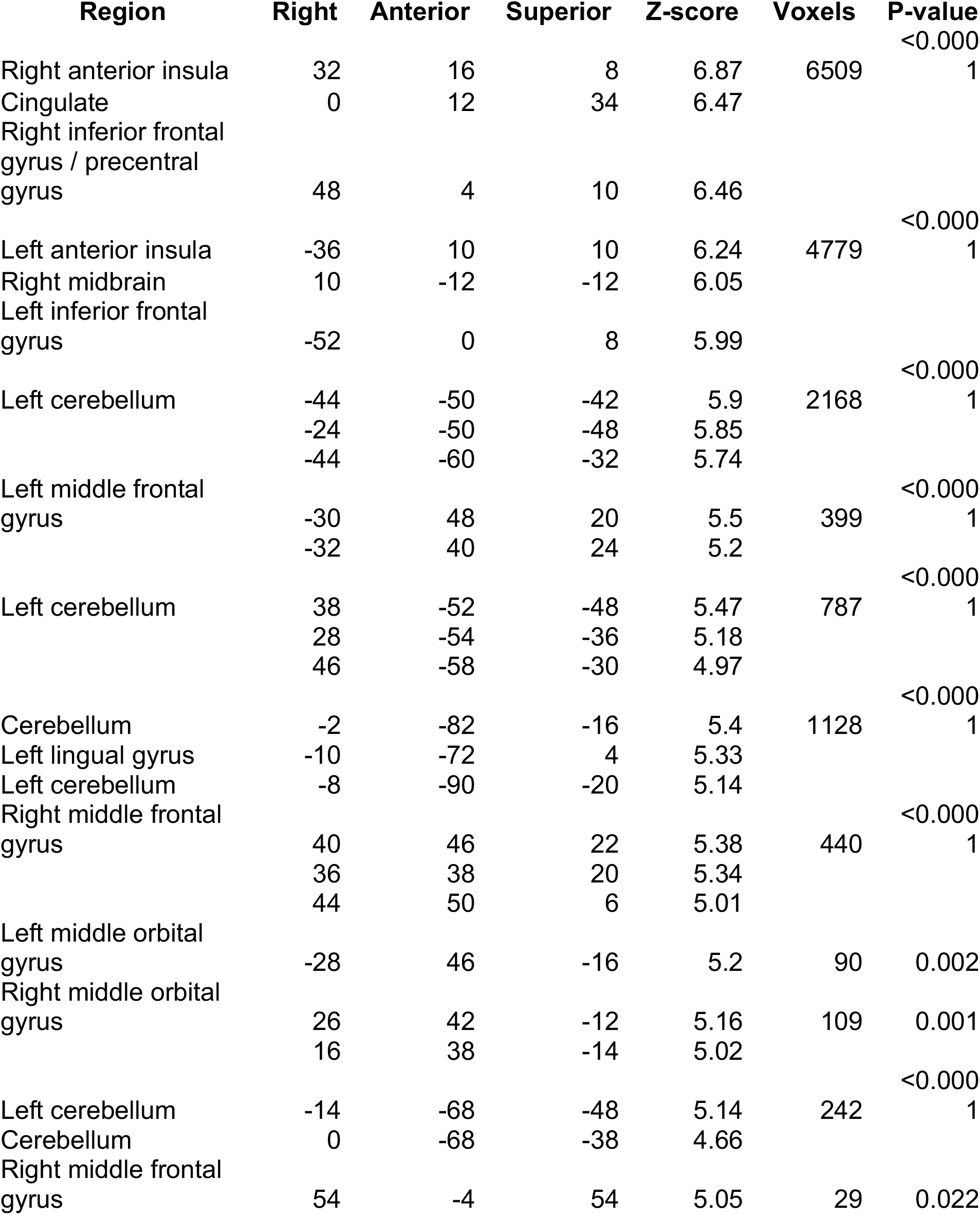
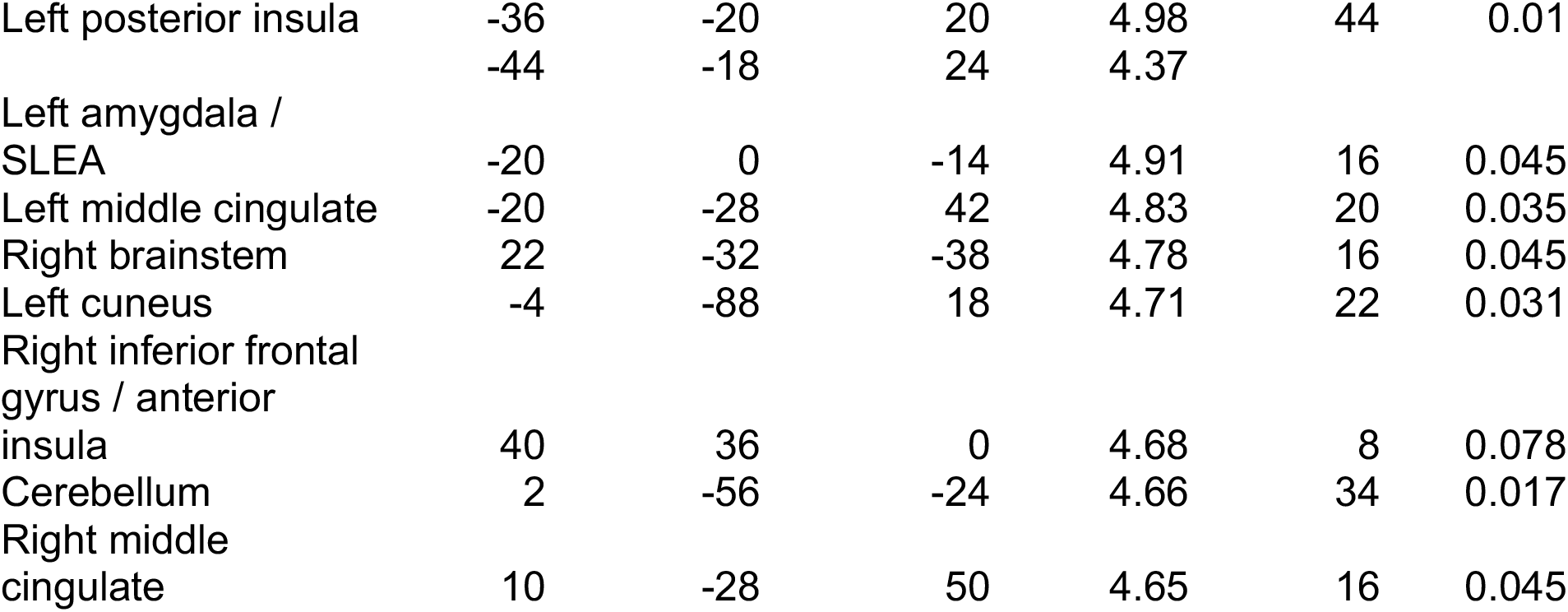
Neural correlates of pain ratings during pain administration in the incidental learning phase, relating to **Fig. 3a.** Initial uncorrected threshold set at p < 0.00001 for interpretable clusters. All p-values are whole-brain FWE-corrected; see also: https://neurovault.org/images/306227/.

**Table 4-1.**
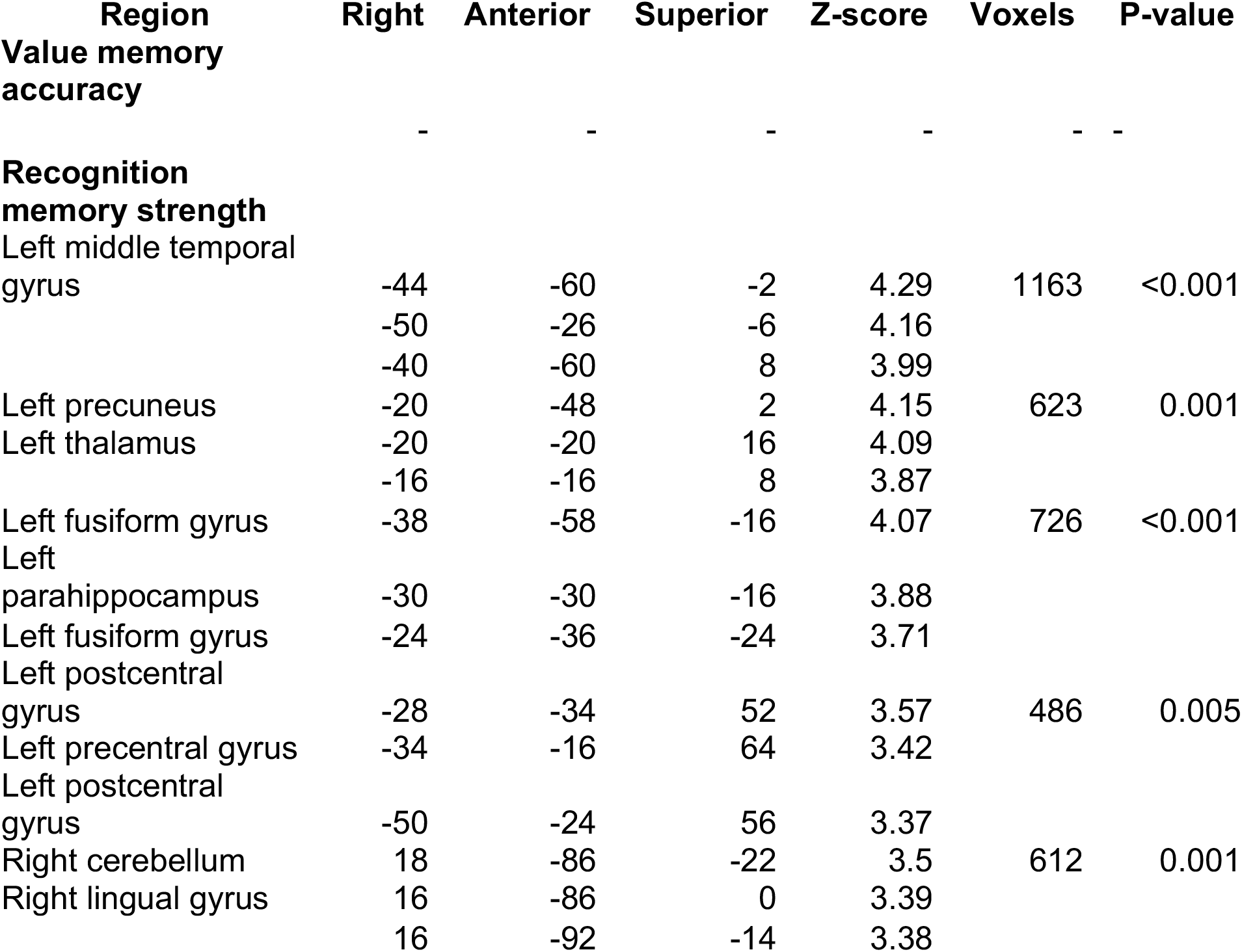
Neural correlates of learning phase subsequent value memory success and subsequent recognition memory strength, related to **Fig. 4c**. All p-values are whole-brain FWE-corrected. For value memory, see: https://neurovault.org/images/504941/; for recognition, see: https://neurovault.org/images/504942/.

**Table 4-2.**
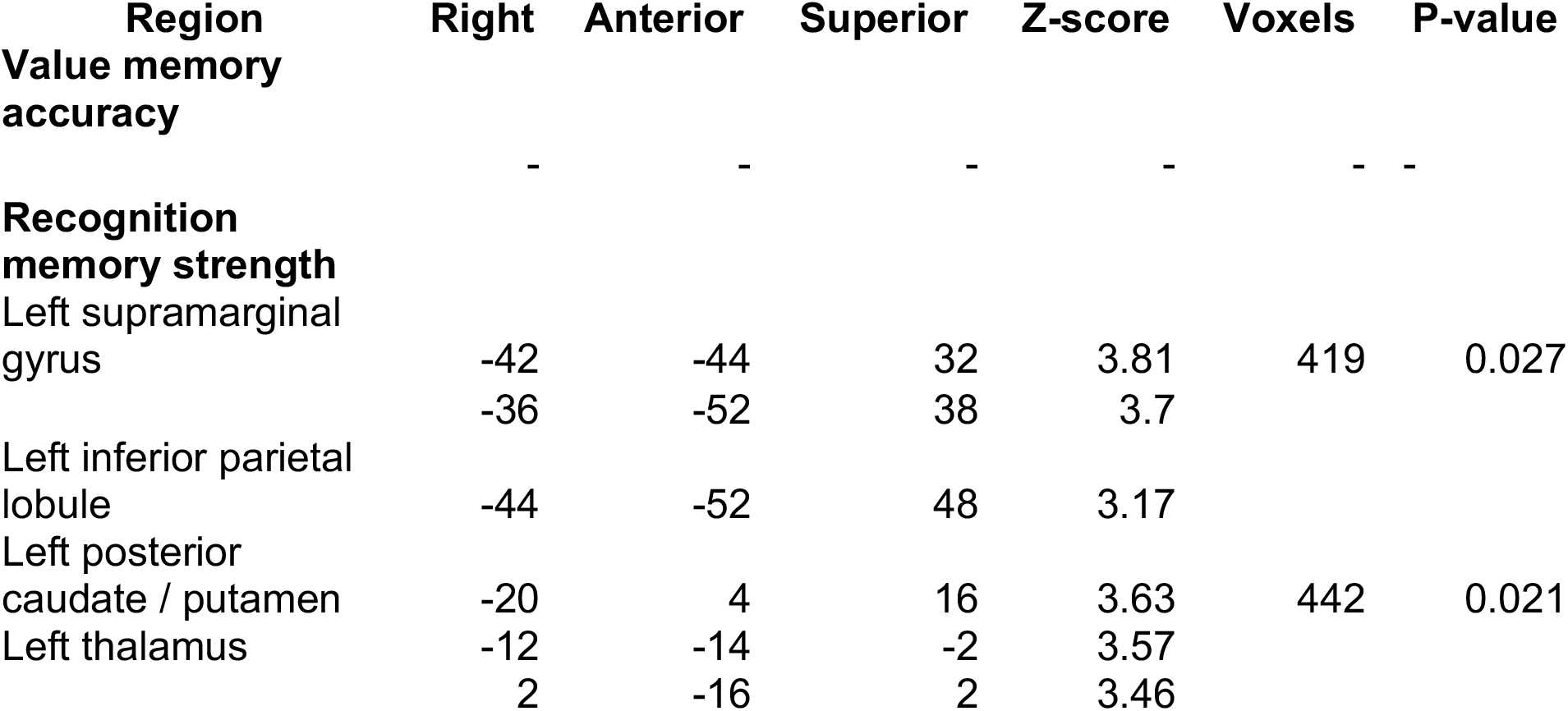
Neural correlates of test phase value memory success and recognition memory strength, relating to **Fig. 4c**. All p-values are whole-brain FWE-corrected. For value memory, see: https://neurovault.org/images/504943/; for recognition, see: https://neurovault.org/images/504944/.

In the PPI analyses, we used the region of the left anterior hippocampus that correlated with behavioral performance as a seed (masking the effect by the hippocampus anatomical mask). The behavioral contrast was trial-by-trial correct versus incorrect later value memory. In the incidental learning phase, we found no overall differences in connectivity for correct versus incorrect later value memory between the anterior hippocampus and any other hippocampal region or brain region. However, individual differences in value memory performance were significantly correlated with the PPI contrast in a region of the left anterior hippocampus (−22, −10, −24; z = 3.78, p = 0.022 SVC; **Fig. 5c**, left) and at a positive but non-significant level in the right anterior hippocampus (22, −12, −22; z = 3.34, p = 0.084 SVC). This correlation indicates that participants with higher value memory performance overall showed greater intra-hippocampal connectivity during successful value memory encoding.

We then conducted a similar connectivity analysis in the memory test phase, again using the left anterior hippocampus as a seed and correct versus incorrect value memory performance as the contrast. We found no overall connectivity differences. Again, however, we found an association between anterior hippocampal connectivity with the bilateral hippocampus and individual differences in value memory performance (left, −22, −10, −22; z = 4.34, p = 0.003 SVC; right, 30 −10 −20; z = 3.69, p = 0.035 SVC; **Fig. 5c**, right). The bilateral hippocampal clusters identified in the learning phase PPI overlapped with the clusters identified in the test phase (**Fig. 5d**). Finally, we examined whether these relationships were selective to value memory performance versus recognition memory. In alternative covariate analyses including recognition memory, we found no significant across-participant correlations in the hippocampus (p-values > 0.73). Together, these results indicate that in individuals with better behavioral discrimination of high versus low heat pain episodes, intra-hippocampal connectivity is stronger during both successful encoding and successful retrieval.

### Control univariate analyses of pain value memory and recognition memory

Finally, while our hypotheses were based on multivariate patterns of activity, for completeness, we also examined the results of several control univariate analyses related to pain, value memory, and recognition memory. First, we looked for overall activation differences at test due to the incidental association of objects with high versus low heat pain, parallel to the multivariate results reported below. We found no evidence of pain value memory reactivation in the test phase in pain-related ROIs, the hippocampus, or across the whole brain (https://neurovault.org/collections/6126/).

Second, we examined univariate correlates of recognition memory strength during learning and test. Note that multivariate analyses analogous to the pain reactivation analyses are not possible with the unbalanced memory strength categories that result from the very high rate of old object recognition (**Fig. 2c**). The univariate models also examined correlates of successful value memory encoding or retrieval, but as noted in the preceding connectivity results, we found no significant results. In the incidental learning phase, we found several clusters at the whole-brain level correlated with recognition memory strength, including the left middle temporal gyrus and left fusiform gyrus (n = 28 participants with memory strength data; **Table 4-1**). Further, we found that activity in the left posterior hippocampus was significantly related to subsequent memory (−18, −42, 4; z = 4.08, p = 0.009 SVC). In a multivariate control analysis, none of these three regions were able to classify pain pattern reactivation at test (p-values > 0.510). In the memory test phase, we found two significant clusters in the left parietal lobe and striatum / thalamus correlated with recognition memory strength at a whole-brain level (**Table 4-2**). Neither of these regions were able to classify pain pattern reactivation at test (p-values > 0.307).

## Discussion

We found that memory for the values of single aversive experiences can guide later decision making and that value memory was related to reactivation of pain-related activity patterns in the hippocampus from the original experience. In our experiments, we first presented incidental, trial-unique object pictures during high or low pain episodes. Then we administered a surprise test where the objects were re-presented in order to measure pain value memory for the incidental pain associations. In a choice experiment, participants successfully avoided pain-associated objects. In an fMRI experiment, pain-related patterns of activity in the hippocampus – but not in traditional pain-associated regions such as the secondary somatosensory cortex and the insula – were reinstated upon re-exposure to objects. Importantly, individual differences in the strength of reactivation in the anterior hippocampus positively related to value memory behavioral performance. Our results suggest that after an aversive experience, a reminder of the event can reactivate neural patterns in the hippocampus to promote avoidance.

The hippocampus is critical for forming memory for episodes, and more generally for forming relational associations between elements of experience (Eichenbaum and Cohen, 2001; Davachi, 2006). Our results suggest that the hippocampus is also important for linking memory for items with representations of value. Importantly, all of our value memory findings were independent of recognition memory strength for the objects. Further, while the heat pain was designed to be salient, and previous studies have reported both pain-related memory improvements and impairments (Schwarze et al., 2012; Forkmann et al., 2013), we found no relationship between recognition memory and either heat pain or pain reactivation. The potential role of the hippocampus in rapid value association learning aligns with research demonstrating that the hippocampus is important for using relational associations to automatically infer reward value (Wimmer and Shohamy, 2012) and to imagine the value of novel experiences (Barron et al., 2013). While these and other studies in humans have primarily shown a role for the hippocampus in reward domains (Lebreton et al., 2009; Peters and Buchel, 2010; Foerde and Shohamy, 2011; Foerde et al., 2013; Gluth et al., 2015; Wimmer et al., 2018), our results suggest that the hippocampus also plays a role in associating episodic experiences with negative value. Our findings additionally accord with the role of the hippocampus in fast learning during contextual fear conditioning in rodents (Phillips and LeDoux, 1994). Building on these findings, an interesting question for future research is whether and how linking an item with value relates to other findings on relational integration (Davachi, 2006; Staresina and Davachi, 2009). Further, a relational account may also predict that the hippocampus is important for encoding associations between experiences and stimulus intensity in general; for example, the coolness of outside temperature, or the intensity of birdsong. However, our association reactivation results involve salient aversive episodic associations, where successful memory can be critical for adaptive behavior.

We found that across participants, stronger reactivation in the anterior hippocampus was related to better value memory performance. The anterior hippocampus has been associated with anxiety (Adhikari et al., 2010; Fanselow and Dong, 2010; Bach et al., 2014) as well as memory integration and generalization (Poppenk et al., 2013; Schlichting et al., 2015; Brunec et al., 2018). Given that episodes of experience never repeat exactly, successful use of previous experiences to guide future decisions is likely to involve significant generalization, which may be facilitated by the anterior hippocampus. Interestingly, in a related study which utilized monetary reward instead of pain, we did not find evidence for hippocampal reactivation of value-related patterns (Wimmer and Buchel, 2016). In addition to the higher salience for pain compared to monetary reward, one difference between these experiments is that the current study utilized a relatively slow trial duration and longer separation of emotional events, potentially leading to better separation of individual episodes and greater hippocampal involvement in memory encoding (Ezzyat and Davachi, 2011).

Recent research has proposed that choices may be guided by sampling representations of previous experiences stored in memory, in contrast to the more common view that choices being are driven by scalar value representations (Hertwig et al., 2004; Lengyel and Dayan, 2005; Weber and Johnson, 2006; Biele et al., 2009; Gluth et al., 2015; Shadlen and Shohamy, 2016). Whether and how agents form successful memory for the value of episodes is a critical component of memory-based models of decision making. Several previous studies provided initial evidence that agents are capable of learning and using the value or “remembered utility” of previous experiences such as freezing cold water or pleasant vacations (Kahneman et al., 1993; Redelmeier and Kahneman, 1996; Fredrickson, 2000; Wirtz et al., 2003). The choice test in our behavioral study, where participants were incentivized to choose the lower-pain object, closely aligns with choice preference measures utilized in classic behavioral economics studies (Fredrickson and Kahneman, 1993; Kahneman et al., 1993). A potential limitation of our fMRI study, however, is that participants made judgments about single items. Nevertheless, a reasonable conservative hypothesis is that the same mechanism supports both retrieval-supported choice behavior and the retrieval of value memory associations for single items.

Memory sampling models of decision making have been supported by recent experimental evidence in humans in the reward domain (Gluth et al., 2015; Murty et al., 2016; Wimmer and Buchel, 2016; Bornstein et al., 2017; Bornstein and Norman, 2017; Enkavi et al., 2017; Bakkour et al., 2019), which our results extend to aversive valuations. Notably, to our knowledge, only the current study and our recent study in the reward domain (Wimmer and Buchel, 2016) examined the influence of episodes in the absence of explicit – and often extensive – participant instructions to remember the features of unique episodes. We focused on incidental encoding, as our goal was to understand the neural mechanisms that may support behavior in non-laboratory environments, where memory formation is often incidental.

From a learning perspective, episodic or single-shot learning of value is not easily accounted for by reinforcement learning models (Sutton and Barto, 1998), even though it is a well-known phenomenon in contextual fear conditioning (e.g. Blanchard et al., 1968) and taste aversion learning (Welzl et al., 2001). Rapid learning can be accomplished in reinforcement learning models with a very high learning rate for positive and negative events, but this leads to memories being erased upon any repetition. One solution involves dynamically decreasing the learning rate based on the number of exposures to a similar situation (e.g. Schiller et al., 2008), which presents a potential convergence with memory sampling accounts. Memory sampling models can naturally implement a decreasing influence of newer experiences: as related episodes accumulate, each new experience added to memory storage will have a lower chance of being sampled and a weaker effect on choice.

Finally, our results provide a novel demonstration that negative emotion-related neural patterns expressed within the same participants during encoding are re-activated at retrieval. Previous studies on memory for negative events (where otherwise neutral stimuli were initially paired with negative content) have either found group-level univariate activation of the same regions across encoding and retrieval or used reverse inference to attribute activation at retrieval to emotion (Maratos et al., 2001; Smith et al., 2004; Erk et al., 2005; Smith et al., 2006; Albanese et al., 2007; Tsukiura and Cabeza, 2008; Kuhl et al., 2010; Fairhurst et al., 2012; Forkmann et al., 2015; Bowen and Kensinger, 2017). In contrast to these univariate approaches, multivariate analyses are more sensitive and directly yield metrics of information content (Poldrack, 2011), while decoding across experiences within-participants removes the need for reverse inference.

We did not find evidence of reactivation of affect-related patterns in value-correlated regions outside of the hippocampus, in contrast to our previous study (Wimmer and Buchel, 2016). Specifically, we found no evidence for the reactivation of pain-related patterns or even univariate activation at test in traditional pain-related regions such as the insula. One potential reason for this null finding could be that in the current design, pain was largely predicted by a preceding cue, resulting in minimal surprise (or prediction error) when heat was administered in conjunction with the incidental object stimuli. This decoupling of the prediction error learning signal from heat onset may have contributed to the relatively lower value memory performance as compared to our previous study using monetary reward (Wimmer and Buchel, 2016). It is possible that in a design in which aversive stimuli are more unexpected, pain-responsive regions may also show reactivation at test. Overall, similarities and differences between memory for positive and negative episode valence are an interesting target for future research.

## Conclusion

Our results demonstrate a mechanism by which memory can support adaptive behavior: patterns of value-related neural activity in the hippocampus from original experiences can be reactivated to guide later decision making. While much is known about how repeated experiences can build simple associations between stimuli and values, the encoding of single episodes has remained relatively unexplored. From everyday experience, it is clear that decisions can be based on single episodes. Given the considerable capacity of episodic memory in humans, memory represents a rich cache of information that can support future decision making. Remembering the value of negative episodes may be particularly important, as avoiding the repetition of highly aversive experiences, such as those involving bodily harm, can help ensure a longer life. Translationally, understanding overactive or underactive reactivation of negative experiences may inform the understanding and treatment of post-traumatic stress disorder, depression, and other mood disorders (Hamilton and Gotlib, 2008; Brewin et al., 2010; Shin and Liberzon, 2010).

## Author Contributions

G.E. Wimmer and C. Büchel designed the experiment; GEW collected and analyzed data; GEW and CB wrote and revised the manuscript.

## Acknowledgments

G.E.W. was supported by ERC-2010-AdG_20100407. C.B. is supported by DFG SFB TRR 289 (Project A02; Project-ID 422744262–TRR 289) and ERC-2019-AdG_883892. We thank Lea Kampermann and Lidia Meißner for assistance with data collection and translation.

## References

Adhikari, A., Topiwala, M.A., and Gordon, J.A. (2010). Synchronized activity between the ventral hippocampus and the medial prefrontal cortex during anxiety. Neuron 65, 257–269.

Albanese, M.C., Duerden, E.G., Rainville, P., and Duncan, G.H. (2007). Memory traces of pain in human cortex. J Neurosci 27, 4612–4620.

Allefeld, C., Gorgen, K., and Haynes, J.D. (2016). Valid population inference for information-based imaging: From the second-level t-test to prevalence inference. Neuroimage 141, 378–392.

Apkarian, A.V., Bushnell, M.C., Treede, R.D., and Zubieta, J.K. (2005). Human brain mechanisms of pain perception and regulation in health and disease. Eur J Pain 9, 463–484.

Atlas, L.Y., Bolger, N., Lindquist, M.A., and Wager, T.D. (2010). Brain mediators of predictive cue effects on perceived pain. J Neurosci 30, 12964–12977.

Bach, D.R., Guitart-Masip, M., Packard, P.A., Miro, J., Falip, M., Fuentemilla, L., and Dolan, R.J. (2014). Human hippocampus arbitrates approach-avoidance conflict. Current biology : CB 24, 541–547.

Bakkour, A., Palombo, D.J., Zylberberg, A., Kang, Y.H., Reid, A., Verfaellie, M., Shadlen, M.N., and Shohamy, D. (2019). The hippocampus supports deliberation during value based decisions. Elife 8.

Barr, D.J., Levy, R., Scheepers, C., and Tily, H.J. (2013). Random effects structure for confirmatory hypothesis testing: Keep it maximal. J Mem Lang 68.

Barron, H.C., Dolan, R.J., and Behrens, T.E. (2013). Online evaluation of novel choices by simultaneous representation of multiple memories. Nat Neurosci 16, 1492–1498.

Biele, G., Erev, I., and Ert, E. (2009). Learning, risk attitudes and hot stoves in restless bandit problems. Journal of mathematical psychology 53, 155–167.

Blanchard, R.J., Dielman, T.E., and Blanchard, D.C. (1968). Prolonged aftereffects of a single foot shock. Psychon Sci 10, 327–328.

Bornstein, A.M., Khaw, M.W., Shohamy, D., and Daw, N.D. (2017). Reminders of past choices bias decisions for reward in humans. Nat Commun 8, 15958.

Bornstein, A.M., and Norman, K.A. (2017). Reinstated episodic context guides sampling-based decisions for reward. Nat Neurosci 20, 997–1003.

Bowen, H.J., and Kensinger, E.A. (2017). Recapitulation of emotional source context during memory retrieval. Cortex 91, 142–156.

Brainard, D.H. (1997). The psychophysics toolbox. Spat Vis 10, 433–436.

Brewin, C.R., Gregory, J.D., Lipton, M., and Burgess, N. (2010). Intrusive images in psychological disorders: Characteristics, neural mechanisms, and treatment implications. Psychol Rev 117, 210–232.

Brooks, J.C., Nurmikko, T.J., Bimson, W.E., Singh, K.D., and Roberts, N. (2002). Fmri of thermal pain: Effects of stimulus laterality and attention. Neuroimage 15, 293–301.

Brunec, I.K., Bellana, B., Ozubko, J.D., Man, V., Robin, J., Liu, Z.X., Grady, C., Rosenbaum, R.S., Winocur, G., Barense, M.D., and Moscovitch, M. (2018). Multiple scales of representation along the hippocampal anteroposterior axis in humans. Current biology : CB 28, 2129–2135 e2126.

Chang, C.C., and Lin, C. (2011). Libsvm: A library for support vector machines. ACM Trans. Intell. Syst. Technol. 2, 1–27.

Cox, R.W. (1996). Afni: Software for analysis and visualization of functional magnetic resonance neuroimages. Comput Biomed Res 29, 162–173.

Davachi, L. (2006). Item, context and relational episodic encoding in humans. Curr Opin Neurobiol 16, 693–700.

Daw, N.D., and Doya, K. (2006). The computational neurobiology of learning and reward. Curr Opin Neurobiol 16, 199–204.

Duncan, K.D., and Shohamy, D. (2016). Memory states influence value-based decisions. J Exp Psychol Gen 145, 1420–1426.

Eichenbaum, H., and Cohen, N.J. (2001). From conditioning to conscious recollection: Memory systems of the brain (New York: Oxford University Press).

Enkavi, A.Z., Weber, B., Zweyer, I., Wagner, J., Elger, C.E., Weber, E.U., and Johnson, E.J. (2017). Evidence for hippocampal dependence of value-based decisions. Sci Rep 7, 17738.

Erk, S., Martin, S., and Walter, H. (2005). Emotional context during encoding of neutral items modulates brain activation not only during encoding but also during recognition. Neuroimage 26, 829–838.

Ezzyat, Y., and Davachi, L. (2011). What constitutes an episode in episodic memory? Psychol Sci 22, 243–252.

Fairhurst, M., Fairhurst, K., Berna, C., and Tracey, I. (2012). An fmri study exploring the overlap and differences between neural representations of physical and recalled pain. PLoS One 7, e48711.

Fanselow, M.S., and Dong, H.W. (2010). Are the dorsal and ventral hippocampus functionally distinct structures? Neuron 65, 7–19.

Fazeli, S., and Buchel, C. (2018). Pain-related expectation and prediction error signals in the anterior insula are not related to aversiveness. J Neurosci 38, 6461–6474.

Foerde, K., Race, E., Verfaellie, M., and Shohamy, D. (2013). A role for the medial temporal lobe in feedback-driven learning: Evidence from amnesia. J Neurosci 33, 5698–5704.

Foerde, K., and Shohamy, D. (2011). Feedback timing modulates brain systems for learning in humans. J Neurosci 31, 13157–13167.

Forkmann, K., Wiech, K., Ritter, C., Sommer, T., Rose, M., and Bingel, U. (2013). Pain-specific modulation of hippocampal activity and functional connectivity during visual encoding. J Neurosci 33, 2571–2581.

Forkmann, K., Wiech, K., Sommer, T., and Bingel, U. (2015). Reinstatement of pain-related brain activation during the recognition of neutral images previously paired with nociceptive stimuli. Pain.

Fredrickson, B.L. (2000). Extracting meaning from past affective experiences: The importance of peaks, ends, and specific emotions. Cognition Emotion 14, 577–606.

Fredrickson, B.L., and Kahneman, D. (1993). Duration neglect in retrospective evaluations of affective episodes. J Pers Soc Psychol 65, 45–55.

Friston, K.J., Worsley, K.J., Frackowiak, S.J., Mazziotta, J.C., and Evans, A.C. (1993). Assessing the significance of focal activations using their spatial extent. Human Brain Mapping 1, 210–220.

Geuter, S., Boll, S., Eippert, F., and Buchel, C. (2017). Functional dissociation of stimulus intensity encoding and predictive coding of pain in the insula. Elife 6.

Gluth, S., Sommer, T., Rieskamp, J., and Buchel, C. (2015). Effective connectivity between hippocampus and ventromedial prefrontal cortex controls preferential choices from memory. Neuron 86, 1078–1090.

Hamilton, J.P., and Gotlib, I.H. (2008). Neural substrates of increased memory sensitivity for negative stimuli in major depression. Biol Psychiatry 63, 1155–1162.

Hebart, M.N., Gorgen, K., and Haynes, J.D. (2014). The decoding toolbox (tdt): A versatile software package for multivariate analyses of functional imaging data. Frontiers in neuroinformatics 8, 88.

Hertwig, R., Barron, G., Weber, E.U., and Erev, I. (2004). Decisions from experience and the effect of rare events in risky choice. Psychol Sci 15, 534–539.

Horing, B., Sprenger, C., and Buchel, C. (2019). The parietal operculum preferentially encodes heat pain and not salience. PLoS Biol 17, e3000205.

Kahneman, D., Fredrickson, B.L., Schreiber, C.A., and Redelmeier, D.A. (1993). When more pain is preferred to less: Adding a better end. Psychol Sci 4, 401–405.

Kriegeskorte, N., Goebel, R., and Bandettini, P. (2006). Information-based functional brain mapping. Proc Natl Acad Sci U S A 103, 3863–3868.

Kuhl, B.A., Shah, A.T., DuBrow, S., and Wagner, A.D. (2010). Resistance to forgetting associated with hippocampus-mediated reactivation during new learning. Nat Neurosci 13, 501–506.

Kurth, F., Zilles, K., Fox, P.T., Laird, A.R., and Eickhoff, S.B. (2010). A link between the systems: Functional differentiation and integration within the human insula revealed by meta-analysis. Brain structure & function 214, 519–534.

Lakens, D. (2017). Equivalence tests: A practical primer for t tests, correlations, and meta-analyses. Soc Psychol Personal Sci 8, 355–362.

Lebreton, M., Jorge, S., Michel, V., Thirion, B., and Pessiglione, M. (2009). An automatic valuation system in the human brain: Evidence from functional neuroimaging. Neuron 64, 431–439.

LeDoux, J.E. (2000). Emotion circuits in the brain. Annu Rev Neurosci 23, 155–184.

Lengyel, M., and Dayan, P. (2005). Hippocampal contributions to control: The third way. In Advances in neural information processing systems 20, J. Platt, D. Koller, Y. Singer, and S. Roweis, eds. (Cambridge: MIT Press), pp. 889–896.

Maratos, E.J., Dolan, R.J., Morris, J.S., Henson, R.N., and Rugg, M.D. (2001). Neural activity associated with episodic memory for emotional context. Neuropsychologia 39, 910–920.

Mazaika, P., Hoeft, F., Glover, G.H., and Reiss, A. (2009). Methods and software for fmri analysis for clinical subjects. In Hum Brain Mapp.

Murty, V.P., FeldmanHall, O., Hunter, L.E., Phelps, E.A., and Davachi, L. (2016). Episodic memories predict adaptive value-based decision-making. J Exp Psychol Gen 145, 548–558.

Peters, J., and Buchel, C. (2010). Episodic future thinking reduces reward delay discounting through an enhancement of prefrontal-mediotemporal interactions. Neuron 66, 138–148.

Phillips, R.G., and LeDoux, J.E. (1994). Lesions of the dorsal hippocampal formation interfere with background but not foreground contextual fear conditioning. Learn Mem 1, 34–44.

Poldrack, R.A. (2011). Inferring mental states from neuroimaging data: From reverse inference to large-scale decoding. Neuron 72, 692–697.

Poppenk, J., Evensmoen, H.R., Moscovitch, M., and Nadel, L. (2013). Long-axis specialization of the human hippocampus. Trends Cogn Sci 17, 230–240.

Rangel, A., Camerer, C., and Montague, P.R. (2008). A framework for studying the neurobiology of value-based decision making. Nat Rev Neurosci 9, 545–556.

Redelmeier, D.A., and Kahneman, D. (1996). Patients’ memories of painful medical treatments: Real-time and retrospective evaluations of two minimally invasive procedures. Pain 66, 3–8.

Rissman, J., and Wagner, A.D. (2012). Distributed representations in memory: Insights from functional brain imaging. Annu Rev Psychol 63, 101–128.

Roy, M., Shohamy, D., Daw, N., Jepma, M., Wimmer, G.E., and Wager, T.D. (2014). Representation of aversive prediction errors in the human periaqueductal gray. Nat Neurosci 17, 1607–1612.

Schiller, D., Levy, I., Niv, Y., LeDoux, J.E., and Phelps, E.A. (2008). From fear to safety and back: Reversal of fear in the human brain. J Neurosci 28, 11517–11525.

Schlichting, M.L., Mumford, J.A., and Preston, A.R. (2015). Learning-related representational changes reveal dissociable integration and separation signatures in the hippocampus and prefrontal cortex. Nat Commun 6, 8151.

Schuirmann, D.J. (1987). A comparison of the two one-sided tests procedure and the power approach for assessing the equivalence of average bioavailability. J Pharmacokinet Biopharm 15, 657–680.

Schultz, W. (2006). Behavioral theories and the neurophysiology of reward. Annu Rev Psychol 57, 87–115.

Schultz, W., Dayan, P., and Montague, P.R. (1997). A neural substrate of prediction and reward. Science 275, 1593–1599.

Schwarze, U., Bingel, U., and Sommer, T. (2012). Event-related nociceptive arousal enhances memory consolidation for neutral scenes. J Neurosci 32, 1481–1487.

Seymour, B., and al., e. (2004). Temporal difference models describe higher-order learning in humans. Nature 429, 664–667.

Seymour, B., O’Doherty, J.P., Dayan, P., Koltzenburg, M., Jones, A.K., Dolan, R.J., Friston, K.J., and Frackowiak, R.S. (2004). Temporal difference models describe higher-order learning in humans. Nature 429, 664–667.

Shadlen, M.N., and Shohamy, D. (2016). Decision making and sequential sampling from memory. Neuron 90, 927–939.

Shin, L.M., and Liberzon, I. (2010). The neurocircuitry of fear, stress, and anxiety disorders. Neuropsychopharmacology 35, 169–191.

Smith, A.P., Henson, R.N., Dolan, R.J., and Rugg, M.D. (2004). Fmri correlates of the episodic retrieval of emotional contexts. Neuroimage 22, 868–878.

Smith, A.P., Stephan, K.E., Rugg, M.D., and Dolan, R.J. (2006). Task and content modulate amygdala-hippocampal connectivity in emotional retrieval. Neuron 49, 631–638.

Staresina, B.P., and Davachi, L. (2009). Mind the gap: Binding experiences across space and time in the human hippocampus. Neuron 63, 267–276.

Stewart, N., Chater, N., and Brown, G.D. (2006). Decision by sampling. Cogn Psychol 53, 1–26.

Sutton, R.S., and Barto, A.G. (1998). Reinforcement learning: An introduction (Cambridge: MIT Press).

Tracey, I., and Mantyh, P.W. (2007). The cerebral signature for pain perception and its modulation. Neuron 55, 377–391.

Tsukiura, T., and Cabeza, R. (2008). Orbitofrontal and hippocampal contributions to memory for face-name associations: The rewarding power of a smile. Neuropsychologia 46, 2310–2319.

Wager, T.D., Atlas, L.Y., Lindquist, M.A., Roy, M., Woo, C.W., and Kross, E. (2013). An fmri-based neurologic signature of physical pain. N Engl J Med 368, 1388–1397.

Weber, E.U., and Johnson, E.J. (2006). Constructing preferences from memory. In The construction of preference, P. Slovic, and S. Lichtenstein, eds. (New York: Cambridge University Press).

Welzl, H., D’Adamo, P., and Lipp, H.P. (2001). Conditioned taste aversion as a learning and memory paradigm. Behav Brain Res 125, 205–213.

Wiech, K., Jbabdi, S., Lin, C.S., Andersson, J., and Tracey, I. (2014). Differential structural and resting state connectivity between insular subdivisions and other pain-related brain regions. Pain 155, 2047–2055.

Wimmer, G.E., and Buchel, C. (2015). Pain to remember: A single incidental association with pain leads to greater memory for neutral items one year later. bioRxiv.

Wimmer, G.E., and Buchel, C. (2016). Reactivation of reward-related patterns from single past episodes supports memory-based decision making. J Neurosci 36, 2868–2880.

Wimmer, G.E., Li, J.K., Gorgolewski, K.J., and Poldrack, R.A. (2018). Reward learning over weeks versus minutes increases the neural representation of value in the human brain. J Neurosci 38, 7649–7666.

Wimmer, G.E., and Shohamy, D. (2012). Preference by association: How memory mechanisms in the hippocampus bias decisions. Science 338, 270–273.

Wirtz, D., Kruger, J., Napa Scollon, C., and Diener, E. (2003). What to do on spring break? The role of predicted, on-line, and remembered experience in future choice. Psychol Sci 14, 520–524.

